# Hypothalamic astrocytes control systemic glucose metabolism and energy balance via regulation of extra-synaptic glutamate signaling

**DOI:** 10.1101/2022.02.16.480737

**Authors:** Daniela Herrera Moro Chao, Matthew K Kirchner, Cuong Pham, Ewout Foppen, Raphael GP Denis, Julien Castel, Chloe Morel, Enrica Montalban, Rim Hassouna, Linh-Chi Bui, Justine Renault, Christine Mouffle, Cristina García-Cáceres, Matthias H. Tschöp, Dongdong Li, Claire Martin, Javier E Stern, Serge H Luquet

**Affiliations:** Université de Paris, CNRS, Unité de Biologie Fonctionnelle et Adaptative, F-75013 Paris, France; Neuroscience Institute, Georgia State University, 30302, Atlanta, Georgia, USA; Center for Neuroinflammation and Cardiovascular Diseases, Georgia State University, 30302, Atlanta, Georgia, USA; Institute of Biology Paris Seine, Neuroscience Paris Seine, CNRS UMR8246, INSERM U1130, Sorbonne Universite, 75005, Paris, France; Helmholtz Diabetes Center (HDC) & German Center for Diabetes Research (DZD), Helmholtz Zentrum München, Neuherberg, 85764, Germany; Division of Metabolic Diseases, Technische Universität München, Munich, 80333, Germany; Medizinische Klinik and Poliklinik IV, Klinikum der Universität, Ludwig-Maximilians-Universität München, Munich, Germany

**Keywords:** astrocyte, PVN, obesity, glucose metabolism, glutamate

## Abstract

The hypothalamus is key in the control of energy balance. However, to this day strategies targeting hypothalamic neurons failed to provide viable option to treat most metabolic diseases. Conversely, the role of astrocytes in systemic metabolic control has remained largely unexplored. Here we show that obesity promotes anatomically restricted remodeling of hypothalamic astrocyte activity. In the paraventricular nucleus (PVN) of the hypothalamus, chemogenetic manipulation of astrocytes results in bidirectional control of neighboring neuron activity, autonomic outflow, glucose metabolism and energy balance. Such process recruits a mechanism involving the astrocytic control of ambient glutamate levels, which becomes defective in obesity. Positive or negative chemogenetic manipulation of PVN astrocyte Ca^2+^ signals respectively worsen or improves metabolic status of diet-induced obese mice. Collectively, these findings highlight a yet unappreciated role for astrocyte in the direct control of systemic metabolism and suggest potential targets for anti-obesity strategy.

## Introduction

Obesity is a public health concern associated with a higher incidence of type 2 diabetes, cardiovascular disease and other life-threatening comorbidities (Morton et al., 2014; Saklayen, 2018) Understanding the mechanisms underlying the onset and progression of the disease is essential to identify potential targets for therapeutic intervention.

Whereas some genetic loci were clearly identified and extensively studied as monogenic causes for obesity, it is widely accepted that the metabolic syndrome is in essence a multifactorial disease that encloses a complex network of molecular, cellular and physiologic alterations. In mammals, energy homeostasis results from the exquisite balance between energy expenditure and nutrient intake. To accomplish this, nervous inputs and circulating factors are integrated by discrete neural circuits in the brain, which, in turn, provide an adaptive behavioral, neuroendocrine and metabolic response (Morton *et al.*, 2014). Over the past 20 years, the hypothalamus has been identified as a major contributor in the pathophysiological processes leading to the metabolic syndrome. (Dietrich and Horvath, 2013). Specialized hypothalamic neuronal populations maintain a strict monitoring of the peripheral metabolic state and regulate food intake and body metabolism by modulating the autonomic nervous system (ANS) (Roh et al., 2016; Timper and Bruning, 2017; Yi et al., 2010). Impairment of appropriate regulation of autonomic outflow has become a key concern in diverse metabolic diseases, including obesity and diabetes (Licht et al., 2013; Thayer et al., 2010). In particular, pre-autonomic neurons located in the paraventricular nucleus (PVN), have emerged as important hypothalamic autonomic/neuroendocrine regulators. They modulate feeding, energy expenditure, neurohormonal response and cardiovascular function, through autonomic efferents projecting to the brainstem and spinal autonomic outputs (Betley et al., 2013; Dampney et al., 2018; Li et al., 2019; Munzberg et al., 2016) and recent genome wide association studies further exemplify that most known and novel obesity-associated gene are concentrated in the hypothalamus (Locke et al., 2015). The activity of hypothalamic neurons is tightly dependent on adequate delivery of energy substrates provided by astrocytes, the major type of glial cells in the brain (Garcia-Caceres et al., 2019; Verkhratsky A, 2018). In physiological and pathological conditions, astrocytes exert a wide spectrum of adaptive functions that can promote and preserve neuronal health, such as synaptogenesis, synaptic efficacy and plasticity within neuronal networks (Freire-Regatillo et al., 2017; Verkhratsky A, 2018). By occupying a strategic position in brain circuits, at the interface between blood vessels and neurons, they participate in the transport and sensing of nutrients and metabolic factors across the blood brain barrier (Allen and Eroglu, 2017).

Recently, hypothalamic astrocytes are being recognized as important players in the control of systemic energy metabolism (Caruso et al., 2013; Garcia-Caceres et al., 2016; Garcia-Caceres et al., 2013; Kim et al., 2014). Astrocytes are impacted by obesity, as central low-grade inflammation triggers inflammatory-like response in hypothalamic astrocytes and induces reactive gliosis in several hypothalamic areas, including the PVN (Buckman et al., 2013; Dalvi et al., 2017; De Souza et al., 2005; Milanski et al., 2009; Robb et al., 2020; Thaler et al., 2012). However, the direct contribution of astrocytes as gate keeper of PVN output, as well as their involvement in the control of body energy and glucose metabolism remain an open question.

In the current study, we took advantages of imaging and gain & loss of function approaches allowing for the direct and selective imaging and manipulation of astrocytes to explore the causal role of astrocyte in metabolic adaptation in physiology and pathophysiological conditions. We first show that while obesity indeed promotes astrocyte calcium (Ca^2+^) activity remodeling, these changes are anatomically defined to hypothalamic sub-nuclei. Among which the PVN showed drastic changes. High fat-induced remodeling of astrocytic network could be mimicked by chemogenetic activation of Gq-coupled Designer Receptors Exclusively Activated by Designer Drugs (DREADDs) while *in vivo* activation or inhibition of PVN astrocyte Ca^2+^signals activity exerts a bidirectional control onto neighboring PVN neuron firing, autonomic outflow, glucose metabolism and energy balance. We found that the underlying mechanism by which astrocyte gate neural activity in the PVN relied on excitatory Amino-Acid Transporters (EAATs)-dependent control of ambient glutamate. In obese mice, this mechanism was selectively impaired and chemogenetic increase (Gq) or decrease (Gi) of astrocytic Ca^2+^ led to metabolic aggravation or amelioration respectively.

In conclusion, our findings show that PVN astrocytes exert a direct control onto systemic glucose metabolism and energy balance and support a concept in which obesity-associated diseases might be at least partially mediated through molecular and signaling modifications in hypothalamic astrocyte-neuron communication.

## Results

### Diet-induced obesity induces anatomically restricted astrocytic Ca^2+^ signals remodeling in the hypothalamus

We first assessed how exposure to energy-rich diet and diet-induced obesity (DIO) alters astrocytic Ca^2+^ signals as a proxy of astrocyte activity (Figure 1A). The genetically encoded Ca^2+^ sensor GCaMP6f was expressed in astrocytes by crossing the tamoxifen-inducible Glast-CreER^T2^ line and the Cre-dependent GCaMP6f line (Pham et al., 2020). Astrocyte-specific expression of GCaMP6f after tamoxifen treatment was evidenced by co-localization of the astrocyte marker s100β and GCaMP6f in stellated cells characteristic of astrocytes (Figure 1B). GCaMP6f-expressing mice were fed either regular diet or high fat high sugar (HFHS, 45% fat and 35% sugar) diet for 2-3-months. Exposure to HFHS diet led to metabolic alteration characterized with increased body weight (Figure 1C), decreased glucose tolerance and increased glucose-induced plasma insulin (**Supplemental Figure 1A, 1B**). Using wide-field fluorescence imaging that enabled the integration of global Ca^2+^ signals, we imaged intrinsic Ca^2+^ activity in hypothalamic astrocytes in acute brain slices. Active regions were identified using the spatio-temporal correlation screening method (Pham *et al.*, 2020) (**Supplemental Figure 1C, 1D**). We observed that basal astrocyte Ca^2+^ signals in obese mice showed an overall increase in the PVN (Figure 1D**, Supplemental Figure 1C, 1D**), the arcuate nucleus (ARC) (Figure 1E) and the dorsomedial nucleus of the hypothalamus (DMH) (**Supplemental Figure 1E**). Interestingly, in the ventromedial nucleus of the hypothalamus (VMH), astrocytes remained relatively protected from diet induced hyperactivity (Figure 1F). Taken together these results indicate that DIO causes heterogeneous and anatomically-restricted remodeling in astrocytic Ca^2+^ activity in metabolically relevant hypothalamic structures.

**Figure 1.**
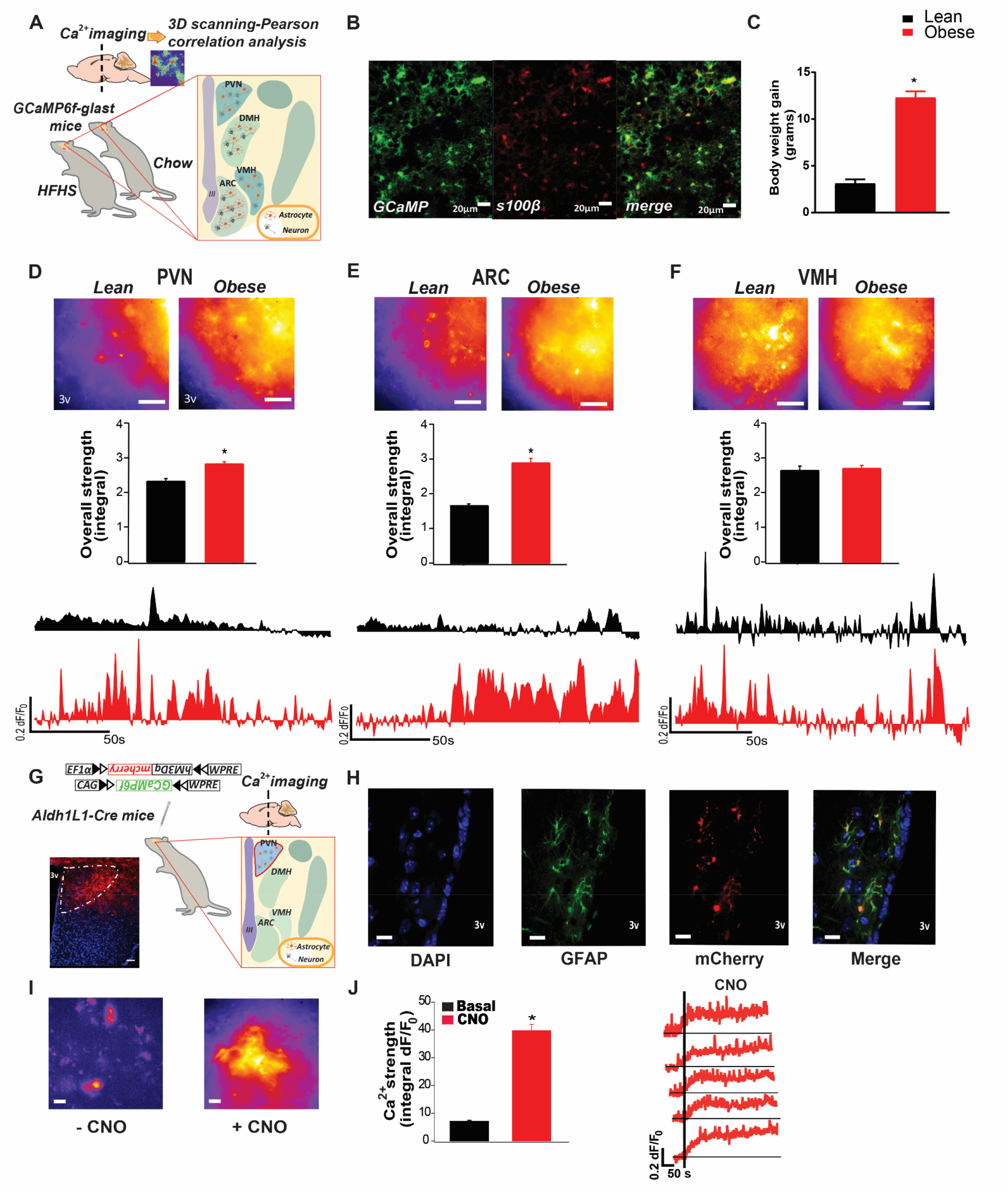
Hypothalamic astrocytes display Ca^2+^ hyperactivity in a heterogeneous manner during diet-induced obesity. **(A)** Experimental setting. **(B)** Representative confocal photomicrograph for GCaMP (green) and s100β (red) immunoreactivity in the PVN of GCaMP6f-Glast-CreER^T2^ mice. **(C)** Body weight gain of GCaMP6f-Glast-CreER^T2^ mice after 4-month chow (lean) or HFHS (obese) diet consumption. **(D-F)** Pseudo-images (top), overall Ca^2+^ strength (middle) and time-course Ca^2+^ signal traces of spontaneous astrocyte activity with under-curve area being shaded (bottom) in **(D)** PVN, **(E)** ARC or **(F)** VMH of lean or obese GCaMP6f-Glast-CreER^T2^ mice. Scale bar: 20µm. **(G)** Experimental setting for viral-mediated approaches in the PVN and representative confocal of corresponding DREADD-mCherry signal (red). Cell nuclei are shown in DAPI (blue), scale bar: 50µm. Co-expression of GCaMP6f and Gq DREADD by dual-AAV injection in the PVN. **(H)** Representative confocal photomicrograph of double immunofluorescence of GFAP (green) and mCherry (red) in the PVN of DREADD injected mice. Cell nuclei are shown in DAPI (blue). Scale bar: 20µm. **(I)** Pseudo-images, **(J)** overall Ca^2+^ strength (left) and representative Ca^2+^ signal traces (right) before and after CNO bath application in PVN astrocytes. Scale bar: 20µm. Histogram data are expressed as mean +/- SEM. See also Figure S1. * P<0.05. For statistical details, see table S1.

### PVN astrocytes control systemic glucose metabolism

Given the prominent role of PVN neural substrate in both autonomic and neuroendocrine control of metabolism (Li *et al.*, 2019; Sutton et al., 2016), we next explored how direct manipulation of PVN astrocyte activity through cell-specific DREADDs engineering affects energy homeostasis. Mice expressing the Cre enzyme under the astrocyte-specific promoter Aldehyde dehydrogenase family 1, member L1 (Cahoy et al., 2008) (Aldh1L1-Cre mice), received an intra-PVN injection of a mixture of Cre-dependent viruses allowing for the expression of (1) mCherry alone (AAV 2/5.EF1α.DIO.mCherry, control) or Gq-coupled hM3Dq DREADD fused with mCherry (AAV 2/5.EF1α.DIO.hM3Dq.mCherry, Gq-DREADD) together with (2) Cre dependent expression of the Ca^2+^ sensor (pAAV5.CAG.Flex.GCaMP6f.WPRE) (Figure 1G). Astrocyte-specific expression of hM3Dq was confirmed by the colocalization of mCherry fluorescent signal with Glial fibrillary acidic protein (GFAP) reactivity (Figure 1H). This combined viral approach allowed to monitor how selective activation of astrocytic Gq coupled hM3Dq by clozapine-N-oxide (CNO) would affect PVN astrocytic intrinsic Ca^2+^ activity in acute brain slices. Compared to minor effect in mCherry expressing control mice (**Supplemental Figure 1F, 1G**), CNO bath application significantly increased astrocytic Ca^2+^ strength in the PVN as a result of hM3Dq activation (Figure 1I, 1J). Using temporal correlation between Ca^2+^ signals as an evaluation of their synchronization, we observed that in contrast to control mice (**Supplemental Figure 1H**), CNO activation of astrocytic hM3Dq drastically increased Ca^2+^ signals synchronization as compared to spontaneous signals (**Supplemental Figure 1I**). This result demonstrates that Gq signaling modulation enables selective activation and synchronization of Ca^2+^ signals in PVN astrocytes.

We next explored how direct manipulation of PVN astrocytes could affect systemic metabolism. Aldh1L1-Cre^+/-^mice received bilateral PVN injection of viruses expressing either mCherry control (AAV5-EF1a-DIO-mCherry; Aldh1L1-Cre^+/-^:PVN^mCherry^) or hM3Dq (AAV5-EF1a-DIO-hM3Dq-mCherry; Aldh1L1-Cre^+/-^:PVN^hM3Dq^) (Figure 2A). To evaluate potential unspecific metabolic consequences of peripheral CNO injections (Gomez et al., 2017), we also injected AAV5-EF1a-DIO-hM3Dq-mCherry in Aldh1L1-Cre^-/-^ mice (Aldh1L1-Cre^-/-^:PVN^hM3Dq^) as a further control. CNO-mediated activation of PVN astrocytes had no consequence on overall whole body energy expenditure (EE), food intake or energy balance in control mice (Aldh1L1-Cre^+/-^ PVN^mCherry^ and Aldh1L1-Cre^-/-^:PVN^hM3Dq^), nor in hM3Dq expressing mice (Aldh1L1-Cre^+/-^ :PVN^hM3Dq^; **Supplemental Figure 2A-I**). However, chemogenetic stimulation of PVN astrocytes by peripheral CNO administration decreased glucose clearance (Figure 2B), increased plasma insulin levels (Figure 2C) and corresponding insulinogenic index (Figure 2D), revealing increased insulin release after an oral glucose tolerance test in Aldh1L1-Cre^+/-^:PVN^hM3Dq^. No significant effect was found in control animals (**Supplemental Figure 2J-Q**).

**Figure 2.**
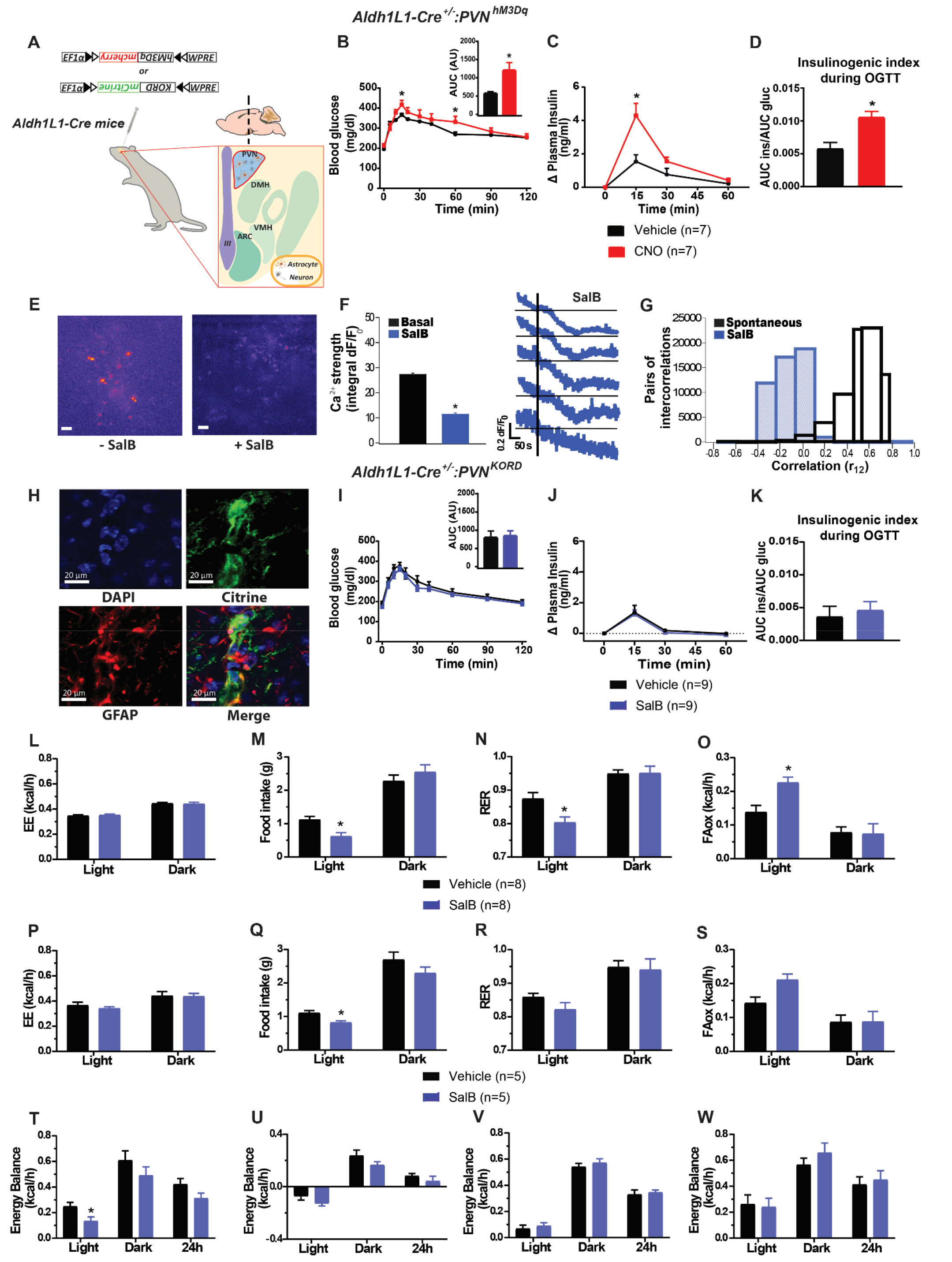
PVN astrocytes regulates glucose metabolism and energy balance. **(A)** Experimental description of viral-mediated approach for Cre-dependent expression of hM3Dq and KORD in the PVN of Aldh1L1-Cre mice. **(B)** Blood glucose and corresponding area under the curve (AUC, top right), **(C)** plasma insulin change and **(D)** insulinogenic index after Vehicle or CNO ip administration followed by an OGTT in Aldh1L1-Cre^+/-^:PVN^hM3Dq^ mice. **(E)** Pseudo-images, **(F)** overall Ca^2+^ strength (left) and time-course Ca^2+^ signal traces (right) of GCaMP signals, **(G)** distribution of temporal correlations of Ca^2+^ responses of all paired active domains (as an estimation of global synchronization) before and after SalB bath application to PVN slices of Aldh1L1-Cre mice expressing KORD receptor. Scale bar: 20µm. Overall Ca^2+^ strength data are expressed as mean +/- SEM. **(H)** Representative confocal photomicrograph of double immunofluorescence of GFAP (red) and mCitrine (green) in the PVN of Aldh1L1-Cre^+/-^:PVN^KORD^ mice. Cell nuclei are shown in DAPI (blue). **(I)**Blood glucose and corresponding AUC (top right), **(J)** plasma insulin change and **(K)** insulinogenic index after Vehicle or SalB ip administration followed by an OGTT in Aldh1L1-Cre^+/-^:PVN^KORD^ mice. **(L-W)** Energy expenditure (EE), food intake, respiratory exchange ratio (RER) and fatty acid oxidation (FAox) after Vehicle or SalB ip injection to **(L-O)** Aldh1L1-Cre^+/-^:PVN^KORD^ and **(P-S)** C57Bl6j:PVN^KORD^ mice. **(T-W)** Energy balance after Vehicle or SalB ip administration in **(T)** Aldh1L1-Cre^+/-^:PVN^KORD^, **(U)** C57Bl6j:PVN^KORD^, **(V)** Aldh1L1-Cre^-/-^:PVN^KORD^ and the control mice **(W)** Aldh1L1-Cre^+/-^ :PVN^mCherry^ mice. Data are expressed as mean +/- SEM. See also Figure S2-4. * P<0.05. For statistical details, see table S1.

Given this striking result, we sought to reproduce this phenomenon in an alternative genetic background. To this aim, C57BL/6 mice received bilateral intra-PVN injection of AAV5-GFAP-hM3Dq-mCherry (C57Bl6j:PVN^hM3Dq^) or control vector AAV5-GFAP-mCherry (C57Bl6j:PVN^mCherry^). While ineffective in control C57Bl6j:PVN^mCherry^ (**Supplemental Figure 3A-D**), CNO-mediated activation of PVN astrocytes in C57Bl6j:PVN^hM3Dq^ resulted in higher glucose-induced insulin release (**Supplemental Figure 3F**), leading to a trend of altered insulin sensitivity as assessed by insulinogenic index (**Supplemental Figure 3G**). These results suggest that chemogenetic manipulation of Gq GPCR pathway in PVN astrocytes readily impairs peripheral glucose metabolism and insulin release, both being the common outcomes preceding insulin resistance states.

We next probed the reciprocal phenomenon by specifically triggering Gi GPCR pathway in PVN astrocytes. Aldh1L1-Cre^+/-^ mice received bilateral injection of AAV5 virus with a modified backbone (pAAV5-EF1a-DIO-HA-KORD-Citrine) to achieve astrocyte, Cre-dependent expression of kappa-opiod receptor (KORD), a recently developed chemogenetic GPCR that can be activated by the pharmacologically inert ligand salvinorin B (SalB) (Vardy et al., 2015) (Figure 2A). After PVN injection of a mixture of virus allowing the co-expression of KORD and GCaMP6f, we could assess astrocyte Ca^2+^ signals before and after KORD activation in acute hypothalamic slices. Bath application of SalB led to a decrease in Ca^2+^ signal strength (Figure 2E, 2F) and global synchronization (Figure 2G) in KORD expressing mice, while these effects appeared marginal in control mCherry expressing mice (**Supplemental Figure 4A-C**). Accordingly, the decrease in intracellular Ca^2+^ release was more pronounced after KORD activation compared to mCherry expressing mice (**Supplemental Figure 4D**). Astrocyte-specific KORD expression was corroborated by the colocalization of mCitrine and GFAP reactivity in PVN astrocytes (Figure 2H).

After validating the cellular consequences of KORD-mediated Gi signaling activation *ex vivo*, we proceeded further *in vivo* to assess the metabolic output arising from decreasing Ca^2+^ activity in vivo in PVN astrocyte. Aldh1L1-Cre^+/-^ (Aldh1L1-Cre^+/-^:PVN^KORD^) or control Aldh1L1-Cre^-/-^ (Aldh1L1-Cre^-/-^:PVN^KORD^) mice received bilateral injections of AAV5-EF1a-DIO-HA.KORD-mCitrine in the PVN. In both control (Aldh1L1-Cre^-/-^:PVN^KORD^) (**Supplemental Figure 4E-L**) and Aldh1L1-Cre^-/+^:PVN^KORD^ mice (Figure 2I-K), peripheral SalB administration did not affect glucose clearance, plasma insulin or insulinogenic index after an oral glucose load.

This result was similar when GFAP-driven expression of hM4Di was achieved in C57Bl6J background (C57Bl6j:PVN^hM4Di^) (**Supplemental Figure 5A-D**). However, given the potency difference between hM4Di and KORD-mediated Gi intracellular signalling (Vardy *et al.*, 2015) and the potential discrepancy between Aldh1l1-CRE and C57Bl6J genetic background, we further explored the metabolic consequence of KORD-mediated Gi signaling in GFAP positive astrocytes. C57Bl6j mice received intra-PVN injection of a mixture of AAV5-GFAP-Cre virus and AAV5-EF1a-DIO-HA.KORD-mCitrine allowing for KORD expression in GFAP-expressing cells upon Cre-mediated recombination (**Supplemental Figure 3I**). KORD expression was visualized by human influenza hemagglutinin (HA) tag presence in GFAP positive astrocytes (**Supplemental Figure 3J**). Here again, Gi signalling initiation through SalB-mediated KORD activation in C57Bl6j:PVN^KORD^ mice did not affect any of the aforementioned parameters (**Supplemental Figure 3K-N**), in line with our observations in Aldh1L1-Cre^+/-^ mice.

Similarly, metabolic efficiency analysis revealed that while peripheral injection of SalB had no effect in either of the control groups (Aldh1L1-Cre^-/-^:PVN^KORD^; Aldh1L1-Cre^+/-^:PVN^mcherry^ **Supplemental Figure 4M-T**), KORD/Gi-mediated modulation of PVN astrocytes did not modify EE (Figure 2L) but induced a decrease in food intake (Figure 2M) and RER (Figure 2N) while increasing FA oxidation (Figure 2O) during the light period. A similar output was observed in C57Bl6j:PVN^KORD^ mice (Figure 2P-S). Accordingly, SalB administration to Aldh1L1-Cre^+/-^ :PVN^KORD^ affected energy balance during the light period (Figure 2T), with a trends for a decrease in C57Bl6j:PVN^KORD^ mice (Figure 2U). No effect of SalB administration was observed in control mice Aldh1L1-Cre^-/-^:PVN^KORD^ and Aldh1L1-Cre^+/-^:PVN^mcherry^ (Figure 2V, 2W). These results suggest that whereas Gq-coupled hM3Dq DREADD modify glucose metabolism and insulin release, activation of astrocytic Gi-KORD receptor in both Aldh1L1 and GFAP expressing astrocytes in the PVN of lean mice promotes only subtle changes in energy utilization.

### PVN astrocytes modulate sympathetic output and thermogenesis

Given the prominent role played by pre-autonomic PVN neurons in autonomic control of energy balance and glucose metabolism (Geerling et al., 2014; Stanley et al., 2019; Stern et al., 2016), we hypothesized that chemogenetic activation of PVN astrocytes could exert a control onto PVN-mediated sympathetic output. As a proxy of autonomic outflow, we first measured change in urine mono-aminergic content as a consequence of chemogenetic activation (Gq) or inhibition (Gi) of PVN astrocytes. ANOVA main effect probability indicated that while no change were observed following CNO or SalB administration in control Aldh1L1-Cre^+/-^:PVN^mCherry^ mice (Figure 3A, 3C), activation of Gq signaling in PVN astrocytes in Aldh1L1-Cre^+/-^:PVN^hM3Dq^ mice significantly increased urine monoamine concentrations (Figure 3B). In contrast, KORD mediated Gi activation showed the opposite effect, with a decrease in urine norepinephrine (NA), as well as tended to decrease dopamine (DA) and Serotonine (5HT) in Aldh1L1-Cre^+/-^:PVN^KORD^ mice (Figure 3D). We further explored if astrocytes in the PVN participate in the regulation of brown adipose tissue (BAT) activity. We monitored *in vivo* BAT metabolic activity by ultrasound and spectral photoacoustic imaging (Clemmensen et al., 2018; Karlas et al., 2019). We observed that PVN astrocyte activation increased local oxygen saturation in BAT blood supply in Aldh1L1-Cre^+/-^PVN^hM3Dq^ mice (Figure 3E, 3F). We next assessed the functional connection between PVN astrocyte activation and thermogenic output. Mice were administered the agonist of cold receptor transient receptor potential cation channel subfamily M member 8 (TRPM8) icilin in order to pharmacologically trigger an adaptive autonomic response similar to the one triggered by cold exposure (Clemmensen *et al.*, 2018). We used BAT monoamine content as a readout of sympathetic outflow (Virtue and Vidal-Puig, 2013). While CNO ip injection in control mice had no significant effect on BAT monoamine content (Figure 3G), astrocyte Gq activation in the PVN promoted increase in adrenaline (AD) and 5HT BAT content in Aldh1L1-Cre^+/^:PVN^hM3Dq^ mice (Figure 3H). Using telemetric core body temperature measurements, we next demonstrated that PVN astrocyte activation through chemogenetic Gq activation was coupled to increased whole body temperature in Aldh1L1-Cre^+/-^:PVN^hM3Dq^ mice (Figure 3I). Finally, we confirmed that Gq signaling activation of PVN GFAP-positive astrocytes in C57Bl6 mice (C57Bl6j:PVN^hM3Dq^; Figure 3J) recapitulated the increase in monoamine content and in BAT (Figure 3K), which were not observed in control C57Bl6j:PVN^mCherry^ mice (Figure 3L). Altogether, these results suggest that PVN astrocytes exert efficient control onto ANS output, thermogenesis and possibly peripheral glucose metabolism.

**Figure 3.**
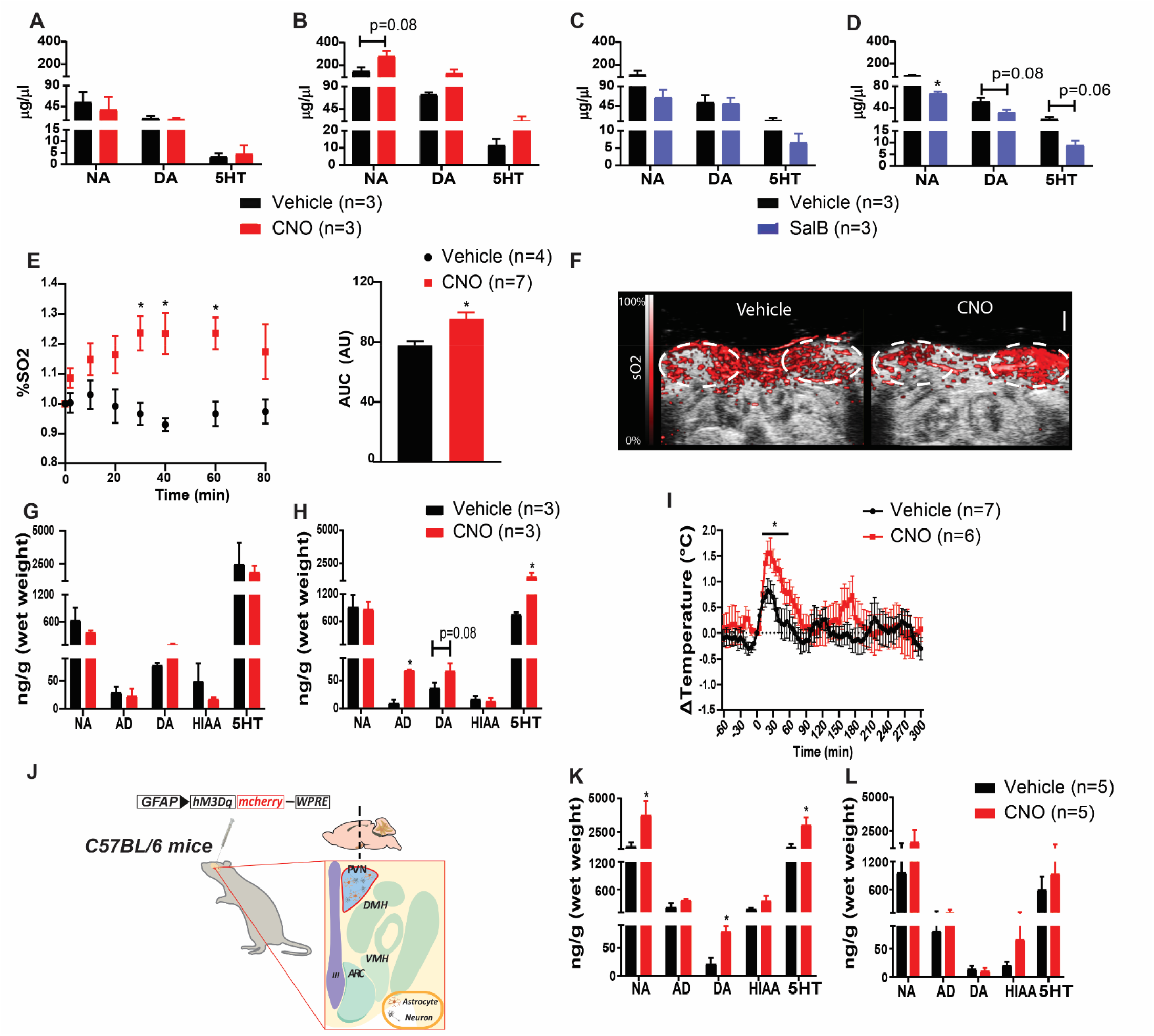
PVN astrocytes regulates sympathetic output and thermogenesis. **(A-D)** Urine mono-aminergic (noradrenaline (NA), dopamine (DA) and serotonin (5HT) concentrations in **(A)** Aldh1L1-Cre^+/-^:PVN^mCherry^ and **(B)** Aldh1L1-Cre^+/-^:PVN^hM3Dq^ mice after Vehicle or CNO ip injection, or in **(C)** Aldh1L1-Cre^+/-^:PVN^mCherry^ and **(D)** Aldh1L1-Cre^+/-^:PVN^KORD^ mice after Vehicle or SalB ip administration. **(E)** Time-course local oxygen consumption (as percentage of blood saturated oxygen (sO_2_)) in BAT (left) and as AUC (right) after Vehicle or CNO ip injection to Aldh1L1-Cre^+/-^:PVN^hM3Dq^ mice. **(F)** Representative images of photoacoustic recording of BAT 40 minutes after Vehicle (left) or CNO (right) ip administration in Aldh1L1-Cre^+/-^:PVN^hM3Dq^ mice. **(G-H)** BAT mono-aminergic (NA, adrenaline (AD), DA, 5-hydroxyindolacetic acid (HIAA) and 5HT content in **(G)** Aldh1L1-Cre^+/-^:PVN^mCherry^ and **(H)** Aldh1L1-Cre^+/-^:PVN^hM3Dq^ mice after Vehicle or CNO ip administration. **(I)** Time-course change in core body temperature of Aldh1L1-Cre^+/-^ :PVN^hM3Dq^ mice after Vehicle or CNO ip injection. **(J)** Representative image of viral delivery of GFAP-hM3Dq in the PVN of C57BL/6 mice. **(K-L)** BAT mono-aminergic (NA, AD, DA, HIAA and 5HT) content in **(K)** C57Bl6j:PVN^hM3Dq^ and **(L)** C57Bl6j:PVN^mCherry^ mice after Vehicle or CNO ip administration. AU refers to arbitrary units. Data are expressed as mean +/- SEM. * P<0.05. For statistical details, see table S1.

### PVN astrocytes exert a bidirectional control on systemic glucose metabolism and energy balance in obese mice

Control C57Bl6j:PVN^mCherry^ mice and C57Bl6j:PVN^hM3Dq^ mice were submitted to 8-weeks of HFHS and assessed for glucose control and metabolic efficiency change induced by PVN astrocyte Gq activation. CNO injection had no effects in obese control mice (**Supplemental Figure 6A-H**), however, in obese C57Bl6j:PVN^hM3Dq^ mice, it led to an acute and drastic deterioration of glucose tolerance (Figure 4A, 4B) associated with large increase in glucose-induced insulin release (Figure 4C), insulinogenic index (Figure 4D) and c-peptide release (Figure 4E) indicative of acute state of insulin-resistance. PVN astrocyte control of glucose metabolism seemed to operate independently from changes in metabolic efficiency, since neither food intake (Figure 4F), RER (**Supplemental Figure 6I**) nor FA oxidation (**Supplemental Figure 6J**) were affected by CNO ip injection to obese C57Bl6j:PVN^hM3Dq^ mice. Whole body energy expenditure was reduced during the dark period (Figure 4G) and accordingly, showed higher positive energy balance (Figure 4H). These results suggest that activation of PVN astrocytes promotes disturbances in systemic glucose metabolism and thereby potentially exacerbates energy storage and metabolic syndrome during obesity.

**Figure 4.**
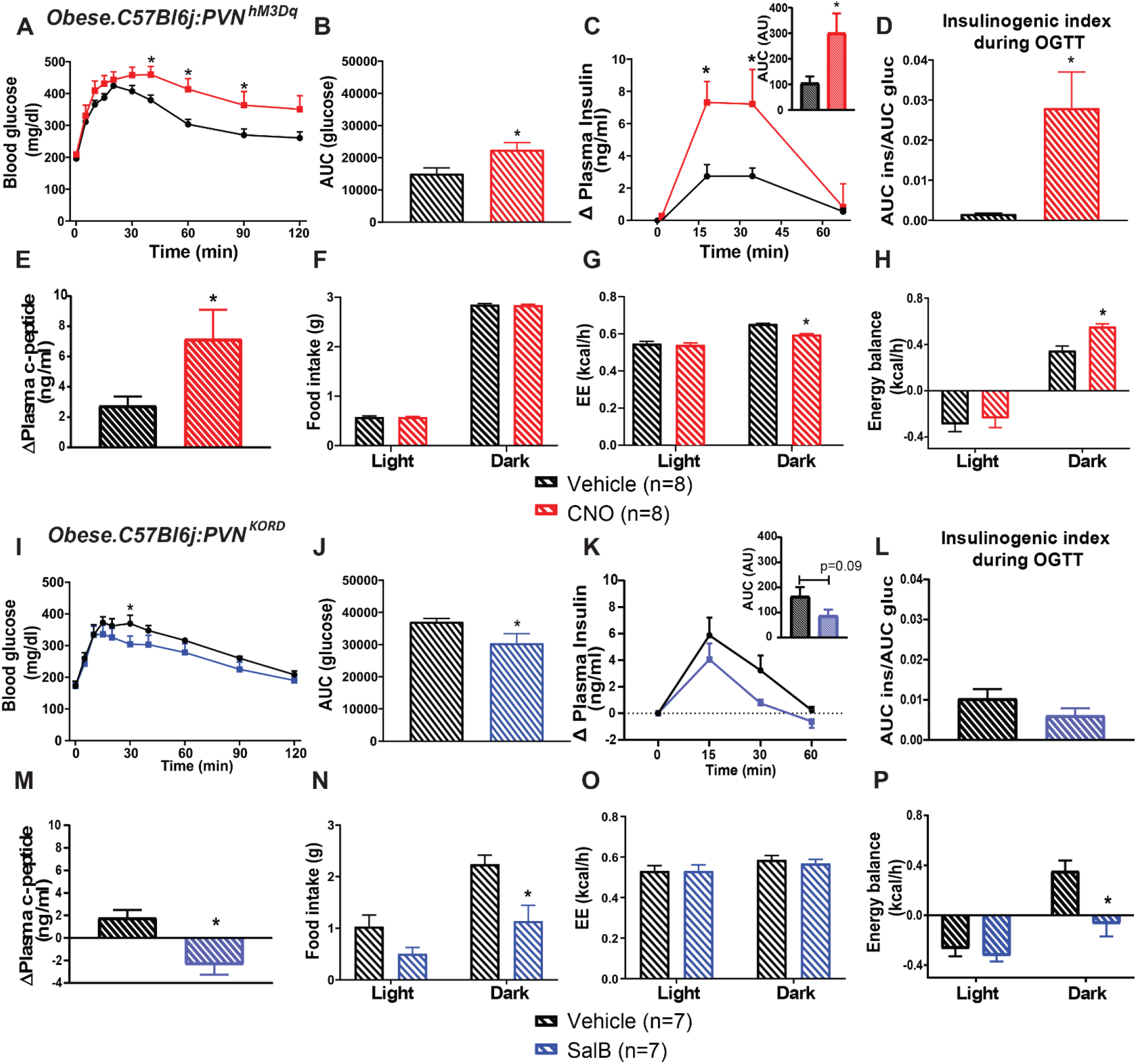
Astrocytes in the PVN exert a bidirectional control on systemic glucose metabolism and energy balance in obese mice. **(A)** Blood glucose, **(B)** glucose AUC, **(C)** plasma insulin change and corresponding AUC (top right), **(D)** insulinogenic index and **(E)** plasma c-peptide change after Vehicle or CNO ip administration followed by an OGTT in C57Bl6j:PVN^hM3Dq^ mice. **(F)** Food intake, **(G)** EE and **(H)** energy balance after Vehicle or CNO ip injection to obese C57Bl6j:PVN^hM3Dq^ mice. **(I)**Blood glucose, **(J)** corresponding AUC, **(K)** plasma insulin change and corresponding AUC (top right), **(L)** insulinogenic index and **(M)** plasma c-peptide change after Vehicle or SalB ip administration followed by an OGTT in obese C57Bl6j:PVN^KORD^ mice. **(N)** Food intake, **(O)** EE and **(P)** energy balance after Vehicle or SalB ip injection to obese C57Bl6j:PVN^KORD^ mice. AU refers to arbitrary units. Data are expressed as mean +/- SEM. See also Figure S5 and Figure S6. * P<0.05. For statistical details, see table S1.

We next probed whether Gi-coupled receptor activation in PVN astrocytes could have opposite effect on metabolic consequence. Gi activation had no impact on glucose tolerance in obese C57Bl6j:PVN^hM4Di^ mice (**Supplemental Figure 5E**), while KORD activation promoted a beneficial effect in obese C57Bl6j:PVN^KORD^ mice (Figure 4I and 4J). In both obese C57Bl6j:PVN^hM4Di^ and C57Bl6j:PVN^KORD^ mice, PVN astrocyte Gi-coupled activation of hM4Di or KORD reduced the level of glucose-induced insulin release during the OGTT (**Supplemental Figure 5F and** Figure 4K), or insulinogenic index (**Supplemental Figure 5G and** Figure 4L). Yet consistently, plasma c-peptide release was reduced following activation of both receptors (**Supplemental Figure 5H and** Figure 4M). Moreover, SalB injection in C57Bl6j:PVN^KORD^ promoted a decrease in food intake (Figure 4N) and RER (**Supplemental Figure 6K**), and an increase in FA oxidation (**Supplemental Figure 6L**), with no changes in energy expenditure (Figure 4O). Finally, contrary to astrocyte Gq receptor activation, KORD-mediated Gi stimulation in PVN astrocytes, promoted negative energy balance during the dark period in obese C57Bl6j:PVN^KORD^ (Figure 4P). These results indicate that PVN astrocytes exert a bidirectional control onto systemic glucose metabolism and energy balance.

### Chemogenetic modulation of astrocyte activity controls PVN parvocellular neuronal firing

We then set out to probe the cellular and molecular mechanism by which PVN astrocytes exert the dynamic control of whole-body metabolic outputs. Parvocellular pre-autonomic neurons in the PVN send efferent projections to autonomic centers in the brainstem and spinal cord to generate essential adaptive metabolic responses for proper systemic glucose metabolism and energy balance regulation (Geerling *et al.*, 2014; Stanley et al., 2010; Stern *et al.*, 2016). We investigated astrocyte-neuron communication by patching parvocellular neurons and recording their activity (Figure. 5A) before and after chemogenetic manipulation of astrocyte Gq or Gi receptor in lean or obese mice. The recorded neurons displayed membrane properties characteristic of parvocellular pre-sympathetic neurons, namely the presence of a low-threshold spike and absence of a transient outward rectification (Luther and Tasker, 2000; Stern, 2001). We combined this approach with simultaneous confocal imaging of astrocyte Ca^2+^ signals after expression of GCaMP6f in either GFAP or Aldh1L1 expressing astrocytes. Activation of Gq-coupled hM3Dq expressing PVN astrocytes by CNO significantly increased neuronal firing frequency from lean C57Bl6j:PVN^hM3Dq^ mice (Figure. 5B, 5C). Conversely, a blunted effect was observed in obese mice (Figure. 5B, 5C). CNO failed to affect firing frequency in the control group of C57Bl6j:PVN^mcherry^ mice (Figure. 5D). We also quantified astrocytic Ca^2+^ activity in an event-based manner using AQuA method (Wang et al., 2019) for both lean and obese C57Bl6j:PVN^hM3Dq^ mice. CNO failed to significantly increase the mean Ca^2+^ event peak in lean or obese C57Bl6j:PVN^hM3Dq^ mice. However, the basal Ca^2+^ peaks in obese mice were significantly higher than lean mice (Figure. 5E, 5F). On the other hand, CNO significantly prolonged Ca^2+^ event duration, and increased Ca^2+^ event synchrony both in lean and obese C57Bl6j:PVN^hM3Dq^ mice (**Figure. 5E, F**). These results are in line with data obtained from wide-field fluorescence imaging (Figure. 1). We then evaluated the latency between astrocytic Ca^2+^ upregulation and the increase in neuronal spike frequency. We found that CNO-induced Ca^2+^ response appeared first in astrocytes followed by neurons, suggesting that changes in astrocyte Ca^2+^ activity significantly preceded those in neuronal firing discharge. These results confirm a sequence of events in which astrocyte Gq activation promotes change in neural activity (Figure. 5G). The latency between activation of astrocytes and upregulation of neuronal spike activity did not differ significantly between lean and obese mice (Figure. 5G**, *right***).

**Figure 5.**
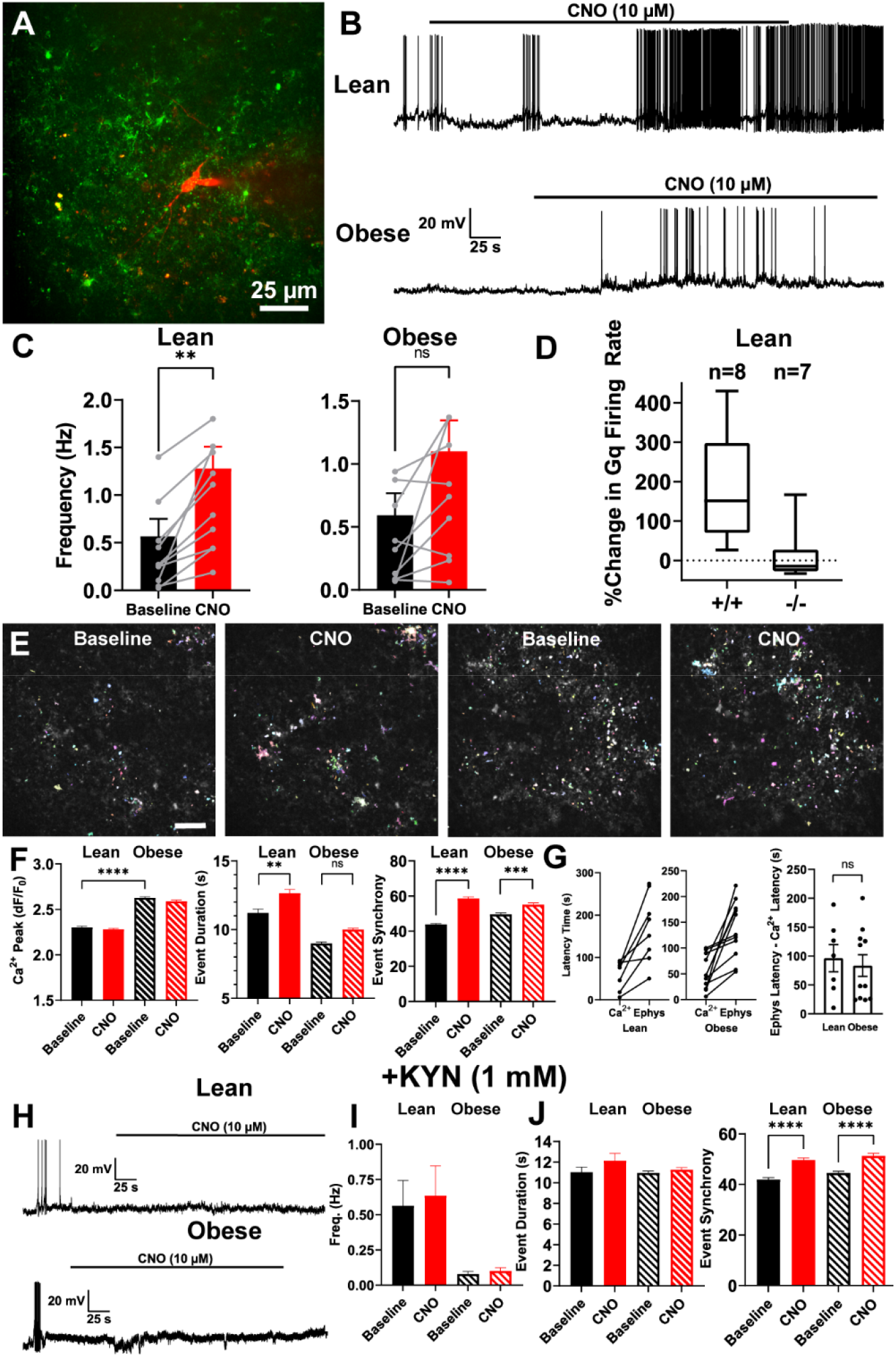
Chemogenetic activation of Gq-coupled receptor in PVN astrocytes enhances parvocellular neuronal firing in lean mice. **(A)** Representative image of a patched parvocellular neuron labelled with Alexa 555 (red) in the PVN of C57Bl6j:PVN^hM3Dq^ mice expressing GCaMP6 (green). **(B)** Firing activity traces in parvocellular PVN neurons in response to CNO in lean (top) and obese (bottom) C57Bl6j:PVN^hM3Dq^ mice. **(C)** Parvocellular PVN neuron spike frequency in response to CNO in lean (left) and obese (right) C57Bl6j:PVN^hM3Dq^ mice. **(D)** %Change in the parvocellular PVN neuron action potential firing frequency after CNO bath application in lean C57Bl6j:PVN^hM3Dq^ (Gq^+/+^) or C57Bl6j:PVN^mCherry^ (Gq^-/-^) mice. **(E)** photomicrograph of Ca^2+^ event analysis overlay in PVN slices of C57Bl6j:PVN^hM3Dq^ mice, in the presence of CNO in both lean (left) and obese (right) animals. Scale bar: 25 μm. **(F)** Summary data of astrocyte Ca^2+^ activity. Ca^2+^ strength (left), event duration (middle) and event synchrony (right) of GCaMP signals in PVN of lean (solid) or obese (stripped) C57Bl6j:PVN^hM3Dq^ mice. **(G)** (Left) Latency of astrocytic Ca^2+^ activity response to CNO application and the upregulation of PVN neuronal spiking activity in lean and obese C57Bl6j:PVN^hM3Dq^ mice. (Right) Difference between the chemogenetic Gq-induced upregulation of Ca^2+^ activity in astrocytes and parvocellular neuronal firing. **(H)** PVN parvocellular firing activity traces recorded from lean (top) and obese (bottom) C57Bl6j:PVN^hM3Dq^ mice, combining bath application of CNO and glutamate channel blocker Kynurenic Acid (KYN, 1 mM). **(I)** Parvocellular PVN neuron spike frequency in the presence of KYN and CNO, in both lean (solid) and obese (stripped) C57Bl6j:PVN^hM3Dq^ mice. **(J)** Summary data of AQuA astrocyte Ca^2+^ activity analysis. Ca^2+^ event duration (left) or event synchrony (right) in the presence of CNO/KYN. Data are expressed as mean +/- SEM. * P<0.05, ** P<0.01, *** P<0.0001. For statistical details, see table S1.

Astrocytes maintain a constant bidirectional communication with neurons, and glutamate is recognized as a key signaling molecule underlying neuro-glial interactions (Durkee and Araque, 2019). Therefore, we investigated if glutamatergic communication is involved in the excitation of parvocellular neurons following astrocyte chemogenetic activation. Blockade of ionotropic glutamate receptors (KYN, 1 mM) prevented the CNO-induced increase in neuronal activity both in lean and obese C57Bl6j:PVN^hM3Dq^ mice (Figure 5H, I). Interestingly, while CNO effects on astrocyte Ca^2+^ event synchrony persisted in the presence of KYN (Figure 5J**, *right***), CNO effects on Ca^2+^ event duration were partially blunted (Figure 5J**, *left***).

We next assessed the effect of astrocyte KORD-Gi-mediated activation on parvocellular neuronal firing. SalB application significantly lowered neuronal activity in acute brain PVN slices of lean Aldh1L1-Cre^+/-^:PVN^KORD^ mice (Figure. 6A, 6B) but had no effect in control Aldh1L1-Cre^+/-^:PVN^mcherry^ mice (Figure. 6C). Importantly, KORD-Gi activation also decreased astrocyte intracellular Ca^2+^ activity, which was manifested as a reduction in event amplitude, event synchrony, and event frequency (Figure. 6D, 6E), while event duration remained unchanged (data not shown). As we observed with chemogenetic activation of Gq in astrocyte, KORD-Gi mediated inhibition of astrocyte Ca^2+^ activity preceded the evoked inhibition of neuronal spike activity (Figure. 6F). Similar to CNO, we found that KYN prevented the SalB-evoked decrease in neuronal activity in Aldh1L1-Cre^+/-^:PVN^KORD^ mice (**Supplemental Figure 7A, 7B**), while the decrease in astrocyte Ca^2+^ activities persisted (**Supplemental Figure 7C**). Taken together, our results indicate that PVN astrocytes exert a bidirectional control over parvocellular firing activity, and that both effects involve a glutamate-dependent mechanism.

**Figure 6.**
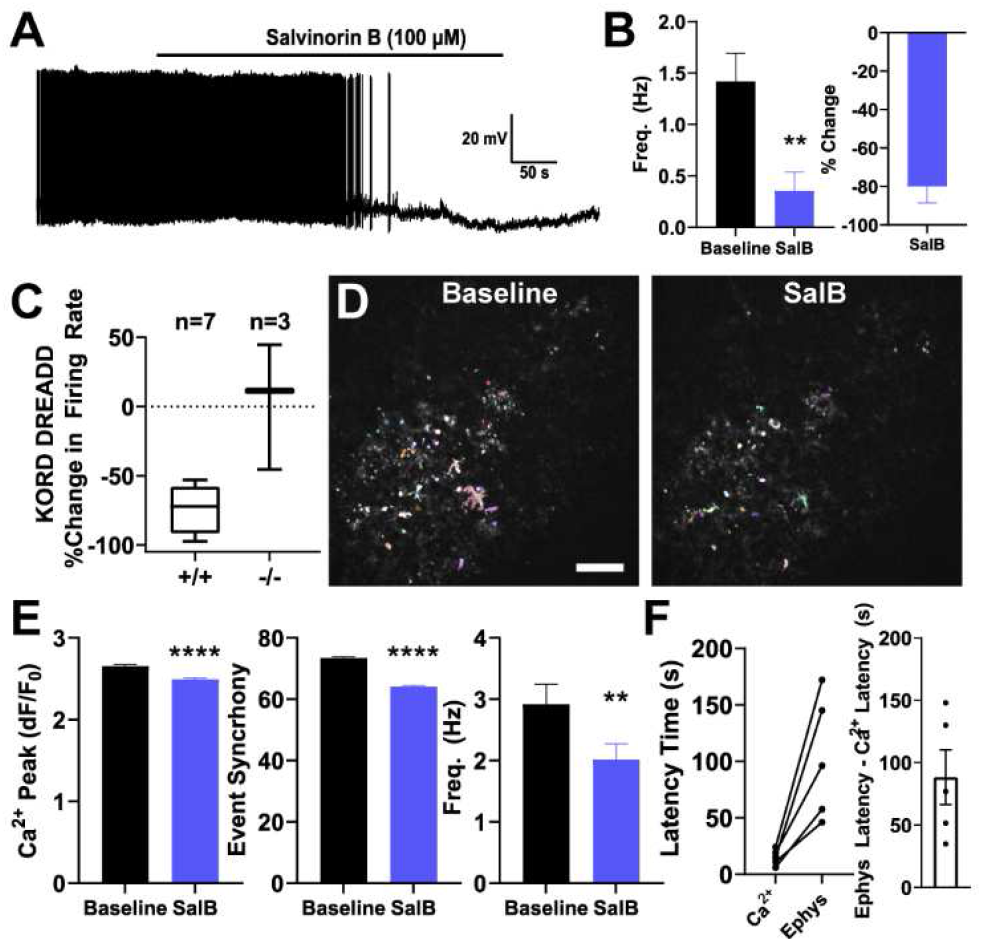
Astrocyte KORD-Gi receptor chemogenetic activation inhibits PVN parvocellular neuronal firing. **(A)** Firing activity traces in parvocellular PVN neurons in response to SalB in Aldh1L1-Cre^+/-^:PVN^KORD^ mice. **(B)** Parvocellular PVN neuron spike frequency (left) and percentage of decrease in firing frequency (right) in the presence of SalB in Aldh1L1-Cre^+/-^:PVN^KORD^ mice. **(C)** %Change in the parvocellular PVN neuron action potential firing frequency after SalB bath application in Aldh1L1-Cre^+/-^:PVN^KORD^ (KORD^+/+^) or Aldh1L1-Cre^+/-^:PVN^mCherry^ (KORD^-/-^) mice. **(D)** Photomicrograph of Ca^2+^ event analysis overlay in PVN slices of Aldh1L1-Cre^+/-^:PVN^KORD^ mice, in the presence of SalB. Scale bar: 25μm. **(E)** Summary data of astrocyte Ca^2+^ activity. Ca^2+^ strength (left), event synchrony (middle) and event frequency (right) of GCaMP signals in PVN of Aldh1L1-Cre^+/-^:PVN^KORD^ mice. **(F)** (Left) Latency of astrocytic Ca^2+^ activity response to SalB application and the upregulation of PVN neuronal spiking activity in Aldh1L1-Cre^+/-^:PVN^KORD^ mice. (Right) Difference between the Gi-induced downregulation of Ca^2+^ activity in astrocytes and parvocellular neuronal firing. Data are expressed as mean +/- SEM. See also Figure S7. * P<0.05, ** P<0.01, *** P<0.0001. For statistical details, see table S1.

### Astrocytic control of extracellular glutamate is impaired during obesity and instrumental in the control of PVN neuron firing

Astrocyte-mediated glutamate signaling to neurons could involve glutamate release and/or modulation of ambient glutamate levels via changes in the activity of astrocyte-selective excitatory Amino-Acid Transporters (EAATs) (Rose et al., 2017). Thus, to determine whether the glutamate-dependent communication between astrocytes and neurons reported above involved modulation of EAAT activity, we repeated experiments in the presence of the glutamate transport blocker DL-*threo*-β-Benzyloxyaspartic acid (TBOA). In this condition, activation of astrocytes by CNO failed to upregulate neuronal activity both in lean and obese C57Bl6j:PVN^hM3Dq^ mice (Figure. 7A, 7C), while CNO effects on astrocyte Ca^2+^ event synchrony persisted (Figure. 7D). Similarly, TBOA also prevented the KORD-mediated inhibition on neuronal firing (**Supplemental Figure 7D, 7E**), without impacting KORD-induced astrocyte Ca^2+^ inhibition (**Supplemental Figure 7F**). Together, these results suggest that the glutamate-dependent increase and decrease of PVN neuronal activity following astrocyte Gq and Gi receptor activation, respectively, is mediated via a bidirectional control of glutamate transporter activity.

**Figure 7.**
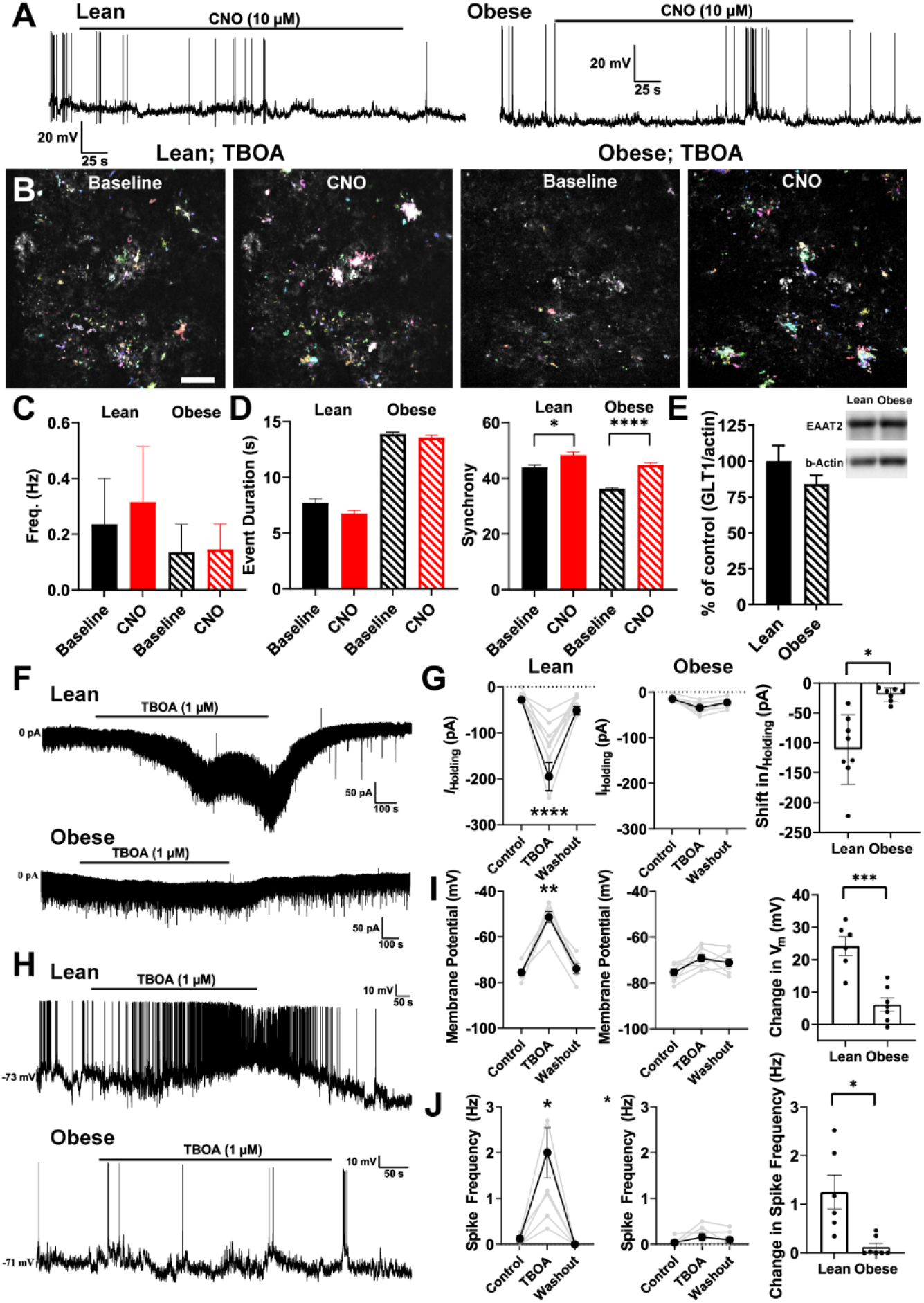
Astrocyte glutamatergic transport signaling contributes to PVN astrocyte-neuron interaction. **(A)** PVN parvocellular firing activity traces recorded from lean (left) and obese (right) C57Bl6j:PVN^hM3Dq^ mice, combining bath application of CNO and glutamate transporter blocker DL-*threo*-β-Benzyloxyaspartic acid (TBOA, 1 μM). **(B)** Representative images of AQuA Ca^2+^ event analysis overlay of GCaMP signals in PVN slices of lean (left) or obese (right) C57Bl6j:PVN^hM3Dq^ mice, demonstrating the strength of Ca^2+^ signals at baseline activity and in the presence of SalB and TBOA. Each image is an average of 10 frames. Scale bar: 25 μm. **(C)** PVN parvocellular neuron spike frequency at baseline activity and in the presence of CNO and TBOA in both lean (solid) and obese (stripped) C57Bl6j:PVN^hM3Dq^ mice. Summary data of astrocyte Ca^2+^ activity in the presence of CNO/TBOA. Ca^2+^ event duration (left) and event synchrony (right) of GCaMP signals in PVN of lean (solid) or obese (stripped) C57Bl6j:PVN^hM3Dq^ mice. **(E)** Protein quantification and representative western blot (inset) of glutamate transporter GLT1 (EAAT2) in lean (solid) and obese (stripped) C57BL/6 mice. **(F)** Membrane and firing properties of parvocellular neurons during voltage clamp, after TBOA (1µM) bath application to PVN slices of lean (top) and obese (bottom) C57BL/6 mice. Holding current (*I*_Holding_). **(G)** Summary graphs representing the shifts in *I*_Holding_ in lean (left), obese (middle), and their difference (right). **(H)** Parvocellular neuron membrane potential (mV) and spike frequency (Hz) in current clamp, during TBOA bath application to PVN slices of lean (top) and obese (bottom) C57BL/6 mice. **(I)** Summary graphs representing the shifts in membrane potential in lean (left), obese (middle), and their difference (right). **(J)** Summary graphs representing the shifts in spike frequency in lean (left), obese (middle), and their difference (right). Data are expressed as mean +/- SEM. * P<0.05, ** P<0.01, *** P<0.0001. For statistical details, see table S1.

Finally, we aimed to determine whether the function and/or expression of astrocyte EEATs is basally altered in obese mice. Western blot analysis of PVN samples showed no significant difference in EAAT2 protein expression levels between lean and obese mice (Figure 7E). We then tested and compared the effects of TBOA *per se* on membrane excitability and firing activity of PVN parvocellular neurons. In voltage clamp mode, we found that bath-applied TBOA produced a significant inward shift in *I*_Holding_ in lean mice (Figure. 7F, 7G), which we previously showed to be due to the slow build-up of glutamate in the extracellular space leading to activation of neuronal extrasynaptic NMDARs (Fleming et al., 2011; Stern *et al.*, 2016; Zhang and Stern, 2017). Surprisingly, TBOA application failed to significantly affect *I*_Holding_ in PVN neurons in obese mice (Figure. 7F, 7G), resulting in a significantly smaller inward shift in *I*_Holding_ in obese compared to lean mice (Figure. 7G**, *right***). Similarly, we found that while TBOA produced a significant membrane depolarization (Figure. 7H, 7I) and increase in spike frequency (Figure. 7J**, *left***) in neurons in lean mice, these effects were absent in obese mice (Figure 7H-J). Taken together, our results showing a blunted effect of TBOA on membrane excitability and firing activity in obese mice support the notion that in this condition, astrocyte glutamate uptake activity is severely blunted. Hence, it is tempting to speculate that obesity-induced alteration of PVN astrocytic glutamate uptake is a leading mechanism contributing to altered astrocyte-neuronal communication and metabolism alteration as a consequence of defective PVN sympathoexcitatory outflow in obese mice.

## Discussion

In the brain, the hypothalamus is regarded as a fundamental player in body weight homeostasis, directly regulating feeding and energy metabolism (Locke *et al.*, 2015; Morton *et al.*, 2014). Perturbations in hypothalamic functions result in obesity and metabolic pathologies (Molavi et al., 2006).

Despite significant endeavor, the approaches that primarily targeted neural substrate have failed to provide successful tools against the metabolic syndrome. This begs for an extended vision of how different brain cell types communicate for the control of energy balance in both physiological and pathophysiological conditions. Indeed, a growing number of studies highlight that glial cells contribute directly to the control of energy homeostasis (Gao et al., 2017; Garcia-Caceres *et al.*, 2016; Kim *et al.*, 2014; Timper et al., 2020; Vicente-Gutierrez et al., 2019; Yang et al., 2015; Zhang et al., 2017) and specific gliopeptide have emerged as critical regulator of the melanocortin-dependant feeding circuit (Bouyakdan et al., 2019). The current vision of how brain cells collaborate in both physiological and pathophysiological conditions have shown the pressing need to understand the glial-neuron communication mechanisms involved in the central regulation of energy balance and to develop anti-obesity approaches that extend beyond primarily targeting neural substrates. In the present study, we mobilized a multidisciplinary set of imaging, genetic and physiologic approaches to demonstrate that hypothalamic astrocytes can causally and directly control whole-body metabolism and the adaptive response to high-fat feeding and obesity in a region-specific manner. We show that PVN astrocytes exert a direct and reversible control on neuron firing, energy balance, systemic glucose metabolism and insulin sensitivity and that specific targeting of astrocyte activity in the PVN exerts beneficial effects on metabolic disturbances during obesity.

Astrocytes represent the most prevalent glial cells in the CNS. The multipartite synapse cradle includes astrocyte and microglial as integral part of synaptic formation (Garcia-Caceres *et al.*, 2019; Semyanov and Verkhratsky, 2021), growth and transmission in multiple regions of the brain (Durkee and Araque, 2019), including the hypothalamus (Bouyakdan *et al.*, 2019; Clasadonte et al., 2017). Astrocytes express a wide arrange of neuropeptide, hormonal and nutrient receptors and exhibit a specific type of excitability mediated by ionic signalling and fluctuations of intracellular Ca^2+^ concentration (Nedergaard et al., 2010). Using Ca^2+^ signals as a proxy of cellular activity, we show that obesity is associated with hyperactivity in hypothalamic astrocytes, recalling the previous association described between high-fat feeding and astrocyte reactivity. In that regard, hypothalamic inflammation and gliosis have been suggested to underlie hypothalamic circuit dysfunction which can lead to hyperphagia, insulin and leptin resistance and consequent obesity (De Souza *et al.*, 2005; Douglass et al., 2017; Reis et al., 2015; Thaler *et al.*, 2012). In addition, abnormal Ca^2+^ glial signaling has been associated to the pathophysiology function of astrocytes and regulation of reactive astrogliosis during brain injury and neuropathology (Denisov et al., 2021; Kanemaru et al., 2013; Nedergaard *et al.*, 2010). We show here a differential susceptibility to obesity-induced astrocyte modifications in the hypothalamus, as obesity affected hypothalamic astrocyte activity in a heterogenous manner, characterized by increased activity in the ARC, DMH and PVN but sparing the VMH. Regionally distinct astrocyte responses have also been observed at different time points after brain injury (Hill et al., 1996), supporting that astrocytes are not a functionally homogeneous cellular population throughout the brain, but rather exhibit inter and intra-regional differences in molecular and cellular profiles (Ben Haim and Rowitch, 2017; Hill *et al.*, 1996; Khakh and Sofroniew, 2015). Given the specificity and ascribed function of hypothalamic nuclei, one can hypothesize that the regionally distinct adaptive response of astrocyte physiology, either beneficial or detrimental to surrounding neurons, will alter feeding, thermogenic adaptation, neuroendocrine and autonomic response through local changes in the ARC, DMH or PVN respectively. In this study we focused our interest on the PVN, given its pivotal role in systemic metabolism control. Indeed, the PVN contains several key neuroendocrine neurons, including oxytocin (OT), vasopressin, corticotropin-releasing hormone (CRH), or thyrotropin-releasing hormone-synthesizing neurons (TRH), in addition to pre-autonomic neurons that project to the dorsal motor nucleus of the vagus nerves (DMX) rostral ventrolateral medulla (RVLM) and the inter-mediolateral cell column (IML), allowing the PVN to directly contribute to both parasympathetic and sympathetic outputs to peripheral tissues (Swanson and Kuypers, 1980; Swanson and Sawchenko, 1980). As an integrative hub, the PVN receives also inputs from the PBN, NTS, DMH and LH (Buijs et al., 2003; Sutton *et al.*, 2016).

Here we found that astrocyte-specific expression and activation of Gq-coupled DREADD receptor in the PVN increases astrocytic Ca^2+^ events (Figure 1 **and Supplemental Figure 1**) as observed following high-fat feeding, while activation of Gi-coupled DREADD led to a decrease of astrocytic Ca^2+^ events and temporal synchronization. Using patch clamp electrophysiology, we show that Gq and Gi DREADD manipulation of PVN astrocytes bidirectionally controlled neighboring parvocellular neuronal firing in a glutamate-dependent manner (Figures 5 and 6). Further, we demonstrate that a core mechanism by which the PVN astrocyte-neuron interaction adapts to nutrient overload involves an adaptive control ambient glutamate levels via changes in the activity of the astrocyte glutamate transporter GLT1 (Figure 7). At the integrated level, chemogenetic-induced Gq or Gi signaling in PVN astrocytes was sufficient to bidirectionally affect circulating and tissue catecholamine content (Figure 3), glucose metabolism and insulin sensitivity *in vivo* (Figure 2 and 4).

Strikingly, manipulating PVN astrocytic Ca^2+^ signals through chemogenetic Gq/Gi engineering respectively aggravated (Gq) or improved (KORD-Gi) the metabolic condition in obese mice. Correspondingly, previous evidence supports beneficial consequences of astrocyte Gi receptor activation, as it corrects astrocyte atrophy and behavioral outputs observed during neurodegenerative diseases (Douglass *et al.*, 2017; Yu et al., 2020). To our knowledge, this is the first demonstration that PVN astrocytes exert a direct and reversible control of energy balance, systemic glucose metabolism and insulin sensitivity. One could even suggest that astrocyte-mediated control of PVN neuron activity might be an essential component of a recently-described functional PVN-pancreas transneuronal circuits directly connecting the PVN to autonomic control beta cell function and glucose metabolism (Papazoglou et al., 2022).

Our data collected using simultaneous *ex-vivo* whole-cell patch clamp electrophysiology and dynamic Ca^2+^ imaging allowed us to establish that chemogenetic control and nutritional modulation of astrocyte networks within the PVN exert a direct and dominant control onto neighboring parvocellular neurons firing activity. Indeed, increases or decreases in astrocyte Ca^2+^ activity achieved through ligand-based activation of Gq or KORD-Gi coupled DREADDs resulted in substantial increases and decreases, respectively, in the firing discharge of PVN neurons. In all cases, changes in astrocytic Ca^2+^ events preceded the one observed in neuronal firing, further establishing that change in PVN neuron firing were a consequence of DREADD-mediated activation of astrocytic Ca^2+^ events.

At the mechanistic level, astrocyte-evoked effects on PVN firing activity were prevented by glutamate receptor blockade, supporting the notion that glutamate signaling is a core component of astrocytic control of PVN neuronal activity. Glutamate-mediated astrocyte-neuron crosstalk could involve active vesicular and non-vesicular glutamate release (Bezzi et al., 1998; Fellin et al., 2004, Araque, 1998 #2; Innocenti et al., 2000). Additionally, astrocytes could also influence ambient levels of glutamate via the activity of potent glutamate transporters, particularly GLT1. Pharmacological block or knockout of the astrocyte GLT1 transporter results in slow buildup of glutamate in the extracellular space, leading to an elevated glutamatergic tone and increased excitability in neighboring neurons (Murphy-Royal et al., 2017; Rothstein et al., 1996).

In our hands, selective blockade of glutamate transport activity prevented the change in neuron firing initiated by either Gq or Gi chemogenetic manipulation of PVN astrocytes (**Supplemental figure 7A, B and 7D, E**) further supporting the notion that astrocyte-neuronal communication critically depended on the control of ambient glutamate levels by the astrocyte glutamate transporter.

Moreover, the delayed firing response of PVN neurons following astrocytic manipulations is consistent with a slow buildup of glutamate and establishment of a glutamatergic tone in the extracellular space. These results align with previous studies pointing at astrocyte GLT1 as a key regulator of astrocyte-neuronal communication in the PVN and supraoptic nucleus (SON) (Fleming *et al.*, 2011; Gordon et al., 2009; Zhang *et al.*, 2017).

In the PVN specifically, angiotensin II activation of Gq AT1 receptors (a phenomenon that is often observed during hypertension) on astrocytes inhibited GLT1 transporter activity, leading to an elevation of extracellular glutamate levels, activation of extrasynaptic N-methyl-D-aspartate receptor (NMDARs) and increased firing of PVN neurons (Stern *et al.*, 2016). Hence, it is possible that Gq DREADD-mediated increase of astrocyte Ca^2+^ activity results in inhibition of GLT1 activity, buildup of extracellular glutamate levels, leading in turn to activation of extrasynaptic NMDARs and subsequent glutamate-mediated increase in PVN firing activity (Fleming *et al.*, 2011; Zhang *et al.*, 2017). Given a similar pharmacological profile and time course of DREADD/Gi-mediated effects, a similar GLT1-based mechanism could mediate the bimodal regulation of PVN neurons by local astrocytes.

Given that 80% of glutamate released undergo active reuptake by astrocyte through GLT-1 and EAAT1 transporters, a critical role for astrocyte glutamate transport in PVN region would have been expected. This is specifically true considering previous observation pointing at glutamate as a primary neurotransmitter mediating body weight regulation by Melanocortin receptor 4 MC4R -bearing neurons in the PVN (Xu et al., 2013). Where our study provides a shift in paradigm is by furnishing experimental evidence that, beyond the buffering glutamate at the multipartite synapse, astrocytes act as gatekeepers of neuronal activity, by being able to establish and impose a degree of freedom through which PVN neurons are allowed to operate. Targeting astrocyte activity, therefore, may offer an upstream control of PVN output, thereby coping with obesity-associated metabolic defect.

We found indeed that obesity is associated with metabolic impairment and disrupted astrocyte-neuronal communication in the PVN. Principally, CNO failed to upregulate neuronal firing in obese mice. This occurred despite activation of DREADDs and modulation of astrocytic Ca^2+^ activity persisted. Thus, we propose that the blunted CNO effect on PVN firing activity is a consequence of a steady-state inhibition of GLT1 activity by basally enhanced astrocyte Ca^2+^ activity in obese mice. This is supported by our results showing an almost completely blunted/occluded effect of TBOA *per se* on the extrasynaptic glutamate tonic current and firing activity of PVN neurons in obese mice. The fact that no changes in GLT1 protein levels were observed further corroborates a blunted activity of GLT1 in astrocytes in this condition. The mechanisms underlying the exacerbated basal astrocyte Ca^2+^ activity and blunted GLT1 activity in obese mice remain at present unknown. Likewise, non-Ca^2+^ dependent mechanism in astrocytes might also contribute to control long-term adaptive change of astrocyte-neurons in response to nutrient overload.

Our data point at extrasynaptic glutamate signaling as being yet an unappreciated mechanism by which astrocytes, at least in the PVN, gate the activity of neighboring neurons. A similar disbalance of extracellular glutamate homeostasis has been commonly observed in many neuropathological processes (Lau et al., 2021; Lewerenz and Maher, 2015; Nedergaard *et al.*, 2010; Stern *et al.*, 2016). Whether these changes in obese mice contribute to exacerbated PVN neuronal activity and sympathoexcitatory outflow from the PVN remains to be determined. Given the pivotal role of the ANS in the control of virtually every metabolically active tissue, a profound mal-adaptation of astrocyte-dependent control of these neurons will inevitably result in a chronic perturbation of peripheral organ activity, a feature that has been proposed as a possible central cause of the metabolism syndrome (Buijs et al., 2006; Marina et al., 2016; Zhang and Stern, 2017). Unfortunately, the tools currently available to finely dissect astrocyte function in vivo with respect to their diversity are still scarce. Hence, we were not able to discriminate whether the entire PVN astrocyte population or a subset of molecularly and anatomically-defined astrocyte subpopulation were involved in systemic metabolic control, neither could we specify the identity of the neurons among the many endocrine and/or pre-autonomic PVN neurons.

In sum, our study demonstrates that PVN astrocyte exert a direct and reversible control of systemic glucose metabolism, insulin sensitivity and energy balance suggesting that future anti-obesity strategies leverage on astrocyte function. Further studies are however warrant to identify the genetic and metabolic signature that specifies sub-population of astrocytes in discrete hypothalamic nuclei. The latter being a requisite to harness the full benefit of astrocyte-directed approaches to cope with metabolic diseases.

## Acknowledgements

This work was funded by the French National Research Agency/Agence Nationale pour la Recherche (ANR) grant # ANR-15-CE14-0030-01, ANR-15-CE14-0030-02 and ANR-15-CE14-0030-03 “Nutripathos”. We acknowledge funding supports from the Centre National de la Recherche Scientifique (CNRS), The Université de Paris and the Foundation pour la Recherche Médicale (FRM). J.E.S was supported from National Heart, Lung, Blood Institute Grant NIH HL090948, National Institute of Neurological Disorders and Stroke Grant NIH NS094640, and funding provided by the Center for Neuroinflammation and Cardiometabolic Diseases (CNCD) at Georgia State University. M.K.K. was supported from National Heart, Lung, Blood Institute Grant F32 HL158172-01. EM was supported by the FRM. CGC was supported from the European Research Council ERC (STG grant AstroNeuroCrosstalk # 757393), the German Research Foundation DFG under Germany’s Excellence Strategy within the framework of the Munich Cluster for Systems Neurology (EXC 2145 SyNergy–ID 390857198) and Helmholtz Excellence Network. MHT was supported from ERC AdG HypoFlam, 695054, European Research Council Executive Agency (ERCEA), DFG Excellence Cluster SyNergy EXC 2145 SyNergy – ID 390857198, German Research Foundation (DFG) and ExNet-0041-Phase2-3, Initiative and Networking Fund of the Helmholtz Association. We thank Giuseppe Gangarossa for scientific and technical expertise. We thank Olja Kacanski for administrative support, Isabelle Le Parco, Ludovic Maingault, Angélique Dauvin, Aurélie Djemat, Magguy Boa and Daniel Quintas for animals’ care and Florianne Michel for genotyping. Telemetry experiments were supported by “The Continuous Glucose Telemetry Award 2018” obtained by Raphaël G.P. Denis and sponsored by Data Sciences International. We acknowledge the technical platforms Functional and Physiological Exploration (FPE) and Bioprofiler of the Université de Paris, CNRS, Unité de Biologie Fonctionnelle et Adaptative, F-75013 Paris, France, the viral production facility of the UMR INSERM 1089 and the animal core facility “Buffon” of the Université de Paris/Institut Jacques Monod. We thank the animal facility of IBPS of Sorbonne Université, Paris. Finally, we would like to thank Xuelong Mi at Virginia Tech for his critical technical support using the AQuA astrocytic calcium analysis.

## Author Contributions

D.H.M.C. and M.K.K contributed to designing research, performing experiments, analyzing data, interpreting results of experiments, preparing figures, and writing of the manuscript. C.P., E.F., R.D., J.C., C.M., E.M., R.H, L.B., J.R., C.M. and O.L., contributed to performing experiments and data analysis. C.G.C. and M.H.T. contributed to funding and critical analysis of experimental plan and conception. D.L, C.M., J.E.S. contributed to conception of the research project, writing of the manuscript, and approving the final version of the manuscript. S.H.L supervised the whole project, secured funding, provided guidance and designed the initial experimental plan and finalized the manuscript with the help of the co-authors.

## STAR Methods

### Resource availability

#### Lead contact and materials availability

- Further information and requests for resources and reagents should be directed to and will be fulfilled by the Lead Contact, Serge Luquet.
- This study did not generate new unique reagents.

#### Data and code availability

- Data reported in this paper will be shared by the lead contact upon request.
- This study did not report original code.
- Any additional information required to reanalyze the data reported in this study is available from the lead contact upon request.

### Experimental models and subject details

#### Animal studies

All animal protocols were approved by the Animal Care Committee of the University of Paris (APAFIS # 2015062611174320), Institut Biologie Paris Seine of Sorbonne University (C75-05-24) or the Georgia State University IAUCUC protocols. Twelve to fifteen-week-old male Aldh1-L1-Cre (Tg(Aldh1l1-cre)) JD1884Htz, Jackson laboratory, Bar Harbor, USA), male C57BL/6J (Janvier, Le Genest St-Isle, France) or male GCaMP6f/Glast-CreER^T2^ (Pham *et al.*, 2020) mice were individually housed at constant temperature (23± 2°C) and submitted to a 12/12h light/dark cycle. All mice had access to regular chow diet (Safe, Augy, France) and water *ad libitum*, unless stated otherwise. Additionally, age matched C57BL/6J or GCaMP6f/Glast-CreER^T2^ mice groups were fed with either chow diet or high-fat high-sugar diet (HFHS, cat n. D12451, Research Diets, New Brunswick, USA) for twelve to sixteen weeks. Body weight was measured every week and body weight gain was estimated as the difference of body weight in week one of HFHS diet consumption to twelve to sixteen weeks after HFHS diet exposure.

### Method details

#### Viral constructs

Designer receptor exclusively activated by designer drugs (DREADD) and GCaMP6f viruses were purchased from http://www.addgene.org/, unless stated otherwise. pAAV-EF1α-DIO-hM3Dq-mCherry (2.4×10^12^ vg/ml, working dilution 1:6, Addgene plasmid #50460-AAV5; http://www.addgene.org/50460/; RRID:Addgene_50460), pAAV-EF1α--DIO-hM4Di-mCherry (2.4×10^12^ vg/ml, working dilution 1:6, Addgene plasmid #50461-AAV5; http://www.addgene.org/50461/; RRID:Addgene_50461), pAAV-EF1α-DIO-mCherry (3.6×10^12^ vg/ml, working dilution 1:4, Addgene plasmid #50462-AAV5; http://www.addgene.org/50462/; RRID:Addgene_50462), pAAV-GFAP-hM3Dq-mCherry (9.1×10^12^ vg/ml, working dilution 1:2, Addgene plasmid #50478-AAV5; http://www.addgene.org/50478/; RRID:Addgene_50478) and pAAV-GFAP-hM4Di-mCherry (3.2×10^11^ vg/ml, working dilution 1:1.5, Addgene plasmid #50479-AAV5; http://www.addgene.org/50479/; RRID:Addgene_50479) were a gift of Bryan Roth. pAAV-GFAP104-mCherry (6.4×10^12^ vg/ml, working dilution 1:1.5, Addgene plasmid #50479-AAV5; http://www.addgene.org/50479/; RRID:Addgene_50479) was a gift of Edward Boyden. pAAV-GFAP-Cre.WPRE (2.2×10^12^ vg/ml, working dilution 1:10, Addgene plasmid #105550-AAV5; http://www.addgene.org/105550/; RRID:Addgene_105550) was a gift from James M Wilson. pAAV-CAG-Flex.GCaMP6f.WPRE (3.15×10^13^ vg/ml, working dilution 1:10, Addgene plasmid #100835-AAV5; http://www.addgene.org/100835/; RRID:Addgene_100835) was a gift of Douglas Kim and GENIE Project. pAAV-GfaACC1D.Lck-GCaMP6f.SV40 (1.53×10^13^ vg/ml, working dilution 1:5, Addgene plasmid #52925-AAV5; http://www.addgene.org/52295/; RRID:Addgene_52925) was a gift of Baljit Khak. In order to produce the pAAV-EF1α-DIO-KORD-mCitrine, the NheI/AscI fragment containing the HA-KORD-Citrine cassette from addgene#65417 was subcloned in Nhe1/AscI backbone pAAV-EF1a-DIO-hM3D(Gq)-mCherry (addgene#50460) to produce pAAV-EF1a-DIO-HA-KORD-Citrine and subsequent adeno-associated virus 5HType 2/5 (2.4×10^12^ vg/ml, working dilution 1:6) were produced by the viral production facility of the UMR INSERM 1089 (Nantes, France).

#### Surgical procedures

For all surgical procedures, mice were rapidly anesthetized with isoflurane (3%), followed by intraperitoneal (ip) injection of analgesic Buprenorphine (Buprecare, 0.3 mg/kg, Recipharm, Lancashire, UK) and Ketoprofen (Ketofen, 10 mg/kg, France) and maintained under 1.5% isoflurane anesthesia throughout the surgery.

##### Stereotaxic surgery

Male Aldh1-L1-Cre^+/-^, Aldh1-L1-Cre^-/-^ and male C57BL/6J mice were placed on a stereotactic frame (David Kopf Instruments, California, USA) and bilateral viral injections were performed (0.3 µl) in the PVN (stereotaxic coordinates: AP −0.8mm, L +0.3mm, V −4.9mm), following the surgical procedures previously described (Berland et al., 2020). Mice recovered for at least 3 weeks after the surgery before being involved in experimental procedures.

##### Continuous body temperature telemetry implantation

Two-weeks after stereotaxic delivery of viral vectors, additional groups of male C57BL/6J mice were implanted with telemetric devices (HD-XG, Data Sciences International (DSI), Minnesota, USA) in the abdominal cavity. After a seven-day period recovery time, the implanted mice were then installed on the DSI receiver. Data were collected using a Ponemah acquisition system (DSI).

#### Photoacoustic BAT imaging

Male Aldh1L1-Cre^+/-^ mice previously injected in the PVN with DREADD viruses were anesthetized under isoflurane and placed on a heated platform. Respiration and cardiac rhythm were constantly monitored. Hair was removed above the interscapular BAT and optic gel was applied before positioning the transducer to ensure a good transmission of the signal. Image acquisition, before and after vehicle (0.9% Sterile sodium chloride solution (Fisher Scientific, Illkrich, France) or Clozapine-N-oxide dihydrocloride (CNO, 0.6mg/kg, Tocris, Bristol, UK) ip injection, was performed using a Vevo LAZR system (FujiFilm VisualSonics, Toronto, ON, Canada) which combine ultrasound and photoacoustic imaging. Light generated from a tunable laser (680–970 nm) is delivered to the tissue through fiber optic bundles integrated into linear-arrays mounted on the ultrasonic transducer (LZ400, fc = 30 MHz). Spectral unmixing analyses were performed with Vevo®Lab software.

#### Indirect calorimetry analysis

After stereotaxic surgery and viral delivery, male Aldh1-L1-Cre^+/-^, Aldh1-L1-Cre^-/-^ and C57BL/6J mice were monitored for metabolic efficiency (Labmaster, TSE Systems GmbH, Bad Homburg, Germany). After an initial period of acclimation in the calorimetry cages, Vehicle (Vehicle matched with CNO injected group: 0.9% Sterile sodium chloride solution (Fisher Scientific, Illkrich, France) or Vehicle matched with SalB injected group: 0.6% DMSO (Sigma-Aldrich, Saint-Louis, USA) in 0.9% sterile sodium chloride solution (Fisher Scientific, Illkrich, France)), CNO (0.6mg/kg dissolved in 0.9% Sterile sodium chloride solution, Tocris, Bristol, UK) or Salvinorin B (SalB, 10mg/kg, dissolved in DMSO and injected in a 0.6% DMSO final solution, Hellobio, Dunshaughlin, Republic of Ireland) were ip injected with a two-day interval between injections depending on the corresponding DREADD expressing group. Food and water intake, whole energy expenditure (EE), oxygen consumption and carbon dioxide production, respiratory exchange ratio (RER=VCO2/VO2, where V is volume) and locomotor activity were recorded as previously described (Berland *et al.*, 2020). Additionally, fatty acid oxidation (Frayn, 1983; Herrera Moro Chao et al., 2019) and energy balance (Hall et al., 2012) were calculated as previously reported. Before and after indirect calorimetry assessment, body mass composition was analyzed using an Echo Medical systems’ EchoMRI (Whole Body Composition Analyzers, EchoMRI, Houston, USA).

#### Oral glucose tolerance test

Viral injected male Aldh1-L1-Cre^+/-^, Aldh1-L1-Cre^-/-^ and C57BL/6J mice and non-injected GCaMP6f/Glast-CreER^T2^ mice were fasted for four hours. A small tail cut incision was performed to take blood samples and measure glycemia with a glucometer (A. Menarini Diagnostics, France). The mice were ip injected firstly with Vehicle, CNO (0.6 mg/kg) or SalB (10 mg/kg) and fifteen minutes after received a glucose oral gavage (2 g/kg). Blood glucose levels were measured after 0, 5, 10, 15, 20, 30, 45, 60, 90 and 120 minutes and blood samples were only taken at 0, 15, 30, and 90 minutes after receiving the glucose oral gavage. Plasma samples were further processed for insulin (mouse ultrasensitive insulin ELISA kit, ALPCO, Salem, USA) and c-peptide (mouse C-peptide ELISA kit, Crystal Chem, IL, USA) measurements. Insulinogenic index (AUC of plasma insulin during OGTT/ AUC of glycaemia during OGTT) was calculated as previously described (Singh and Saxena, 2010).

#### *Ex-vivo* epifluorescence calcium imaging

Male Aldh1-L1-Cre^+/-^ mice previously injected with GCamP6f and DREADDs viral constructs and GCaMP6f/Glast-CreER^T2^ mice were terminally anaesthetized using isoflurane. Brains were removed and placed in ice-cold oxygenated slicing artificial cerebrospinal solution (aCSF, 30mM NaCl, 4.5mM KCl, 1.2mM NaH_2_PO_4_, 1mM MgCl_2_, 26mM NaHCO_3_, and 10mM D-Glucose and 194mM Sucrose) and subsequently cut into 300-µm thick PVN coronal slices using a vibratome (Leica VT1200S, Nussloch, Germany). Next, brain slices were recovered in aCSF (124mM NaCl, 4.5mM KCl, 1.2mM NaH_2_PO_4_, 1mM MgCl_2_, 2mM CaCl_2_, 26mM NaHCO_3_, and 10mM D-Glucose) at 37 °C for 60 minutes. Imaging was carried out at room temperature under constant perfusion (∼3 ml/min) of oxygenated aCSF. The overall cellular fluorescence of astrocytes expressing GCaMP6f was collected by epifluorescence illumination. A narrow-band monochromator light source (Polychrome II, TILL Photonics, Germany) was directly coupled to the imaging objective via an optical fiber. Fluorescence signal was collected with a 40x 0.8NA or a 63x 1.0NA water immersion objective (Zeiss, Germany) and a digital electron-multiplying charge-coupled device (EMCCD Cascade 512B, Photometrics, Birmingham, UK) as previously described (Pham, 2020). A double-band dichroic/filter set was used to reflect the excitation wavelength (470 nm) to slices and filter the emitted GCaMP6 green fluorescence (Di03-R488/561-t3; FF01-523/610, Semrock). The same filter was used for slices expressing both GCaMP6 and DREADD-mCherry. PVN slices were transferred to the imaging chamber, where 3-minute astrocyte spontaneous activity recordings were performed in slices of GCaMP6f/Glast-CreER^T2^ mice. In the case of PVN slices of Aldh1-L1-Cre^+/-^ mice, we performed a basal epifluorescence recording (60 seconds), followed by a 120 second bath application of CNO (10µM) or SalB (100µM) and 240 seconds recording over the washing of the compounds.

The responsive regions displaying Ca^2+^ signals were scrutinized by the three-dimensional spatio-temporal correlation screening method (Pham *et al.*, 2020). Background signal was subtracted from the raw images by using the minimal intensity projection of the entire stack. Ca^2+^ signals of individual responsive regions were normalized as dF/F_0_, with F_0_ representing the baseline intensity and quantified using Matlab (The MathWorks, France) and Igor Pro (Wavemetrics, USA). We gauged signal strength of Ca^2+^ traces of single responsive regions by calculating their temporal integration and normalizing per minute. The global temporal synchronization of detected Ca^2+^ signals was determined by the temporal Pearson’s correlation coefficients of all combinations between single Ca^2+^ regions (Pham *et al.*, 2020).

#### Simultaneous *ex-vivo* whole-cell patch clamp electrophysiology and confocal Ca^2+^ imaging

Viral injected male Aldh1-L1-Cre^+/-^ and C57BL/6J mice were anesthetized with pentobarbital (50 mg kg^-1^ ip) and subsequently perfused through the heart with 30 mL of ice cold aCSF sucrose solution with NaCl replaced by equal-osmol sucrose (in mM: 200 sucrose, 2.5 KCl, 1 MgSO_4_, 26 NaHCO_3_, 1,25 NaH_2_PO_4_, 20 D-Glucose, 0.4 ascorbic acid, and 2.0 CaCl_2_; pH 7.2; 300-305 mosmol l^-1^). The brain was subsequently removed, mounted in the chamber of a vibratome (Leica VT1200s, Leica Microsystems, Buffalo Grove, IL, USA) using superglue and the ventral surface pressed firmly against a block of 3% KCl agar. The brain was submerged in sucrose solution and bubbled constantly with 95% O_2_/5% CO_2_. Coronal slices (240 µm thickness) containing the PVN were cut and placed in a holding chamber filled with aCSF and bubbled with 95% O_2_/5% CO_2_. The aCSF is identical in composition to the sucrose solution, but with 200 mM sucrose replaced by 119 mM NaCl. The slice chamber was warmed using a water bath at 32°C for 20 minutes before placement at room temperature for a total minimum of 60 minutes rest before proceeding with the experiment.

Slices were placed into a chamber on the stage of a microscope and perfused constantly (∼3 ml/min) with aCSF bubbled continuously with 95% O_2_/5% CO_2_ and warmed to 32°C. Parvocellular neurons were targeted for patch clamp using morphological properties such small soma size relative to their magnocellular counterparts, as well as their proximity to as many GCaMP-expressing astrocytes as possible. Whole cell current clamp recordings were obtained from PVN parvocellular neurons using pipettes (2.5-4 MΩ) pulled from borosilicate glass (o.d. 1.5 mm) using a P-97 flaming/brown horizontal micropipette puller (Sutter Instruments, Novato, CA). The pipette internal solution consisted of (in mM): 135 KMeSO_4_, 8 NaCl, 10 HEPES, 2 Mg-ATP, 0.3 Na-GTP, 6 phosphocreatine, 0.2 EGTA with pH 7.2-7.3 and 285-295 mOsmol (kg H_2_O)^-1^. The liquid junction potential for the KMeSO_4_ internal was approximately −10 mV and was not corrected. Occasionally, Alexa 555 (50 μM, Invitrogen, MA, USA) was included to visualize the neuron. For current clamp recordings, traces were obtained with an Axopatch 200B amplifier (Axon Instruments, Foster City, CA) and digitized using an Axon 1440B Digitizer (Axon Instruments, Foster City, CA) at 10 kHz on a desktop computer running Clampex 10 software (Molecular Devices). In the experiments where DL-threo-beta-Benzyloxyaspartate (TBOA, 1 µM, Tocris, Bristol, UK) alone was bath applied, cells were recorded in voltage clamp, held at −70 mV, and filtered at 2 kHz. Data were discarded if series resistance exceeded a 20% change over the course of the recording. To positively confirm a patched cell was indeed a parvocellular neuron, a series of increasingly negative current injections was applied to the cell from a resting membrane potential of −60mV. The activation of a low threshold depolarization in response to the cessation of current injection confirms parvocellular identity (Tasker and Dudek, 1991). Basal recordings of parvocellular and astrocytic Ca^2+^ activity were firstly performed for a minimum of 250 seconds before bath application of either CNO (10 µM) or SalB (100 µM) for 250 seconds. Additional recordings adding CNO or SalB combined with kynurenic Acid (KYN, 1 mM, Abcam, Cambridge, UK) or TBOA (1 µM, Tocris, Bristol, UK), were performed after washout for a minimum of 5 minutes after initial application of CNO/SalB. Data were analyzed using either Clampfit or Igor Pro 8 (Wavemetrics Inc.)

Simultaneously, GCaMP6f expression in astrocytes was visualized using the Dragonfly 200 laser spinning disk confocal imaging system and an iXon 888 EMCCD camera (Andor Technology, Belfast, UK). Using Andor’s FUSION software, we captured time series images with z-stacking to maximize acquisition of astrocytic Ca^2+^ events near the patched neuron. To quantify network-wide Ca^2+^signals within the confocal plane, we employed the Astrocyte Quantification and Analysis (AQuA) method that extracts event-based information from optical sectioning image series (Wang *et al.*, 2019). Image background was subtracted, and images were converted from FUSION’s native file format to TIFFs in ImageJ (NIH). Imaging videos of astrocytic GCaMP were then processed using the AQuA script run through Matlab (v2019b) (MathWorks, Natick, MA, USA). Full details of the script functionality and code can be found at: https://github.com/yu-lab-vt/AQuA. The parameters used for the analysis are as follows: intensity threshold scaling factor 1, smoothing (sigma) 1.5, minimum Size (pixels) 24, temporal cut threshold 2, growing z threshold 1, rising time uncertainty 2, slowest delay in propagation 2, propagation smoothness 1, z score threshold 2, maximum distance 0, minimum correlation 0, maximum time difference 2, temporally extended events disabled, ignore delay tau enabled. We filtered out events lasting longer than 60 seconds and whose amplitude exceeded a 10 dF/F_0_ change in amplitude to minimize inclusion of artifacts in the analysis. Event data were imported into Prism 8 for analysis (GraphPad, San Diego, CA, USA). Some waveform data and figures were generated in Igor Pro (Wavemetrics, Portland OR, USA).

#### Brain tissue Immunofluorescence

Viral injected male Aldh1-L1-Cre^+/-^ and C57BL/6J mice received Vehicle, CNO or SalB ip injection. Sixty minutes later, the mice were euthanized with pentobarbital (500 mg/kg, Dolethal, Vetoquinol, France). An additional group of GCaMP6f/Glast-CreER^T2^ mice was also euthanized. Mice were transcardially perfused with 0.1 M sodium phosphate buffer (PBS, pH 7.5) followed by 4% paraformaldehyde in phosphate buffer (0.1 M, pH 7.2). Brains were removed and post-fixed overnight in 4% paraformaldehyde. Afterwards, the brains were transferred to 30% sucrose in PBS for 2 days for cryoprotection. Next, 25 µm brain sections were cut in a freezing cryostat (Leica, Wetzlar, Germany) and further processed for immunofluorescence following the procedure previously described (Berland *et al.*, 2020). Free-floating brain sections were incubated at 4°C overnight with chicken anti-Green fluorescent protein (GFP, 1:500, Aves Labs, Davis, California, USA), rabbit anti-s100β (1:500, Abcam, Cambridge, UK), rabbit Ds Red (1:1000, Living colors, Takara Bio, St-Germain-en-Laye, France), mouse anti-Glial fibrillary acidic protein (GFAP, 1:1000, Sigma-Aldrich, Saint-Louis, USA) or rabbit anti-HA epitope tag (1:2000, Rockland, Limerick, USA) primary antibodies. The next day, sections were rinsed in Tris-buffered saline (TBS, 0.25M Tris and 0.5M NaCl, pH 7.5) and incubated for 2 hours with secondary antibodies (1:1000, Thermo fisher Scientific, MA, USA) conjugated with fluorescent dyes: goat anti-chicken Alexa 488, donkey anti-rabbit Alexa 594, donkey anti-mouse Alexa 488 and donkey anti-rabbit Alexa 647. After rinsing, the sections were mounted and coverslipped with DAPI (Vectashield, Burlingade, California, USA) and examined with a confocal laser scanning microscope (Zeiss LSM 510, Oberkochen, Germany) with a color digital camera and AxioVision 3.0 imaging software.

#### Western Blotting

C57BL/6J mice were euthanized with pentobarbital (500 mg/kg, Dolethal, Vetoquinol, France) and shortly after decapitated. The mouse head was then immediately immersed in liquid nitrogen for 4 seconds. After removal of the brain, the anterior hypothalamus was dissected on ice-cold surface and brain lysates were prepared in a solution containing 1% Sodium dodecyl sulfate (SDS, Sigma-Aldrich, Saint-Louis, USA), 0.2% phosphatase inhibitors (phosphatase inhibitor cocktail, Sigma-Aldrich, Saint-Louis, USA) and 0.1% protease inhibitors (protease inhibitor cocktail, Roche, Boulogne-Billancourt, France). Protein content in the hypothalamic lysates was measured with a BC Assay protein quantification Kit (Interchim Uptima, Montlucon, France). Western blotting was performed as previously described (Berland *et al.*, 2020). Membranes were immunoblotted with rabbit anti-EEAT2 (1:500, Novus Biologicals, Abingdon, UK). HRP-coupled secondary antibody anti-rabbit (1:10000, Cell Signaling Technology, Charles Renard, France) was matched with an ECL detection system. Quantification was performed using Image J software (NIH).

#### Mono-aminergic content determination

Viral injected male Aldh1-L1-Cre^+/-^ and C57BL/6J mice were injected ip with Vehicle or CNO, followed by icilin ip (5 mg/kg, Tocris, Bristol, UK) and sacrificed with pentobarbital (500 mg/kg, Dolethal, Vetoquinol, France) after sixty minutes. Interscapular brown adipose tissue (BAT) was dissected and immediately snap frozen in liquid nitrogen. Additional viral injected male Aldh1-L1-Cre^+/-^ and C57BL/6J mice received Vehicle, CNO or SalB ip injection and were euthanized sixty minutes later. Urine (removed from the bladder) was collected and immediately stored at −80°C. Monoamine analyses were performed at the Bioprofiler platform of the Unit “Biologie Fonctionnelle et Adaptative”, University of Paris, BFA, UMR 8251 CNRS, F75205, Paris, France. Catecholamines and metabolites were analyzed by reversed phase-High-performance Liquid Chromatography using a Shimadzu system connected to a Waters 2465 electrochemical detector (HPLC-ED, Waters, USA) with a glassy carbon working electrode (0.7V, 10nA). BAT was homogenized in an ice-cold solution, containing 0.4% ethylenediaminetetraacetic acid and 0.1M perchloric acid, while urine samples were processed using MonoSpin PBA columns (Interchim, Montlucon, France). After centrifugation, the supernatant was further analyzed with HLPC-ED. Mobile phase (58,5 mM Sodium Acetate (Sigma, St Louis, USA), 0.7mM Octane Sulfonic Acid (Sigma O 0133, St Louis, USA), pH 3,8 for mobile phase A and 100% MeOH for mobile phase B) was pumped at a flow rate of 1ml/min and monoamines and metabolite concentrations were detected at an oxidation potential of 750mV compared to the reference electrode. Compounds were separated by an isocratic flow (93% A / 10% B) using a Kromasil® AIT column (length 250mm; internal diameter 4.6mm; particle size 5µm) at 30°C. Catecholamines and metabolites were quantified using LabSolution software (Shimadzu, Kyoto, Japan) by integration of the peak absorbance area, employing a calibration curve established with known catecholamine concentrations.

### Quantification and statistical analysis

#### Statistical analysis

Results are expressed as the mean ± SEM. All statistical analysis was performed using GraphPad Prism 8 (La Jolla, CA, USA). Data were analyzed by T-test, two-way or one-way ANOVA depending on each experimental design. Pairwise comparisons were evaluated with a Tukey post-hoc test. Blood glucose, plasma insulin, %SO_2_ and body temperature values were analyzed by ANOVA of repeated measures with 2 between subject factors: *Time* and *Substance*. An adjustment of Bonferroni was performed with significant values set as p<0.05. All details related to statistical analyses are summarized in Table S1.

