## Supplemental information for "Hypothalamic astrocytes control systemic glucose metabolism and energy balance via regulation of extra-synaptic glutamate signaling"

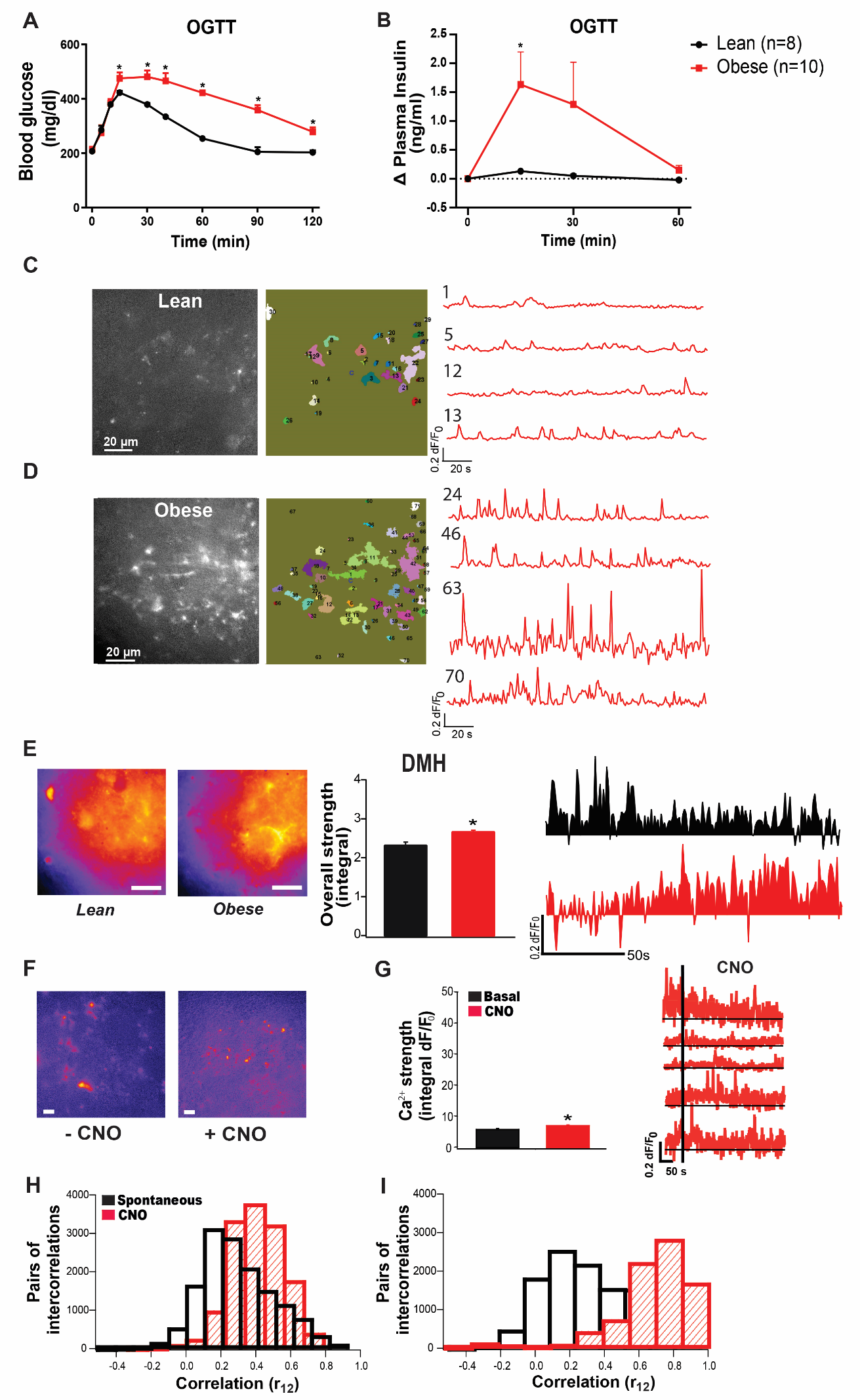

**Supplementary Figure 1. (A)** Blood glucose and **(B)** plasma insulin change during an OGTT in GCaMP6f-Glast-CreER^T2^ mice after 4-month chow (lean) or HFHS (obese) diet consumption. **(C-D)** Average projection of time-lapse GCaMP epifluorescence images (left), detected Ca^2+^ active domains (middle) and representative signal traces (right) of spontaneous Ca^2+^ activity in the PVN of **(C)** lean or **(D)** obese mice. **(E)** Fluorescence projection of global GCaMP6 signals from time-lapse images of DMH of lean or obese GCaMP6f-Glast-CreER^T2^ mice (left), comparison of overall Ca^2+^ signal strength (middle) and representative spontaneous Ca^2+^ signal with under-curve area being shaded in. Scale bar: 20µm. **(F)** Fluorescence projection, **(G)** overall Ca^2+^ strength (left) and time-course Ca^2+^ signal traces (right) of GCaMP signals in the PVN of Aldh1L1-Cre mice expressing mCherry control viral construct before and after CNO bath application. Scale bar: 20µm. Histogram data are expressed as mean +/- SEM. **(H-I)** Distribution of temporal correlations between Ca^2+^ responses of all paired active domains (as an estimation of global synchronization) before and after CNO bath application to PVN slices of Aldh1L1-Cre mice expressing **(H)** hM3Dq receptor or **(I)** mCherry control viral construct.* P<0.05. For statistical details, see table S1.

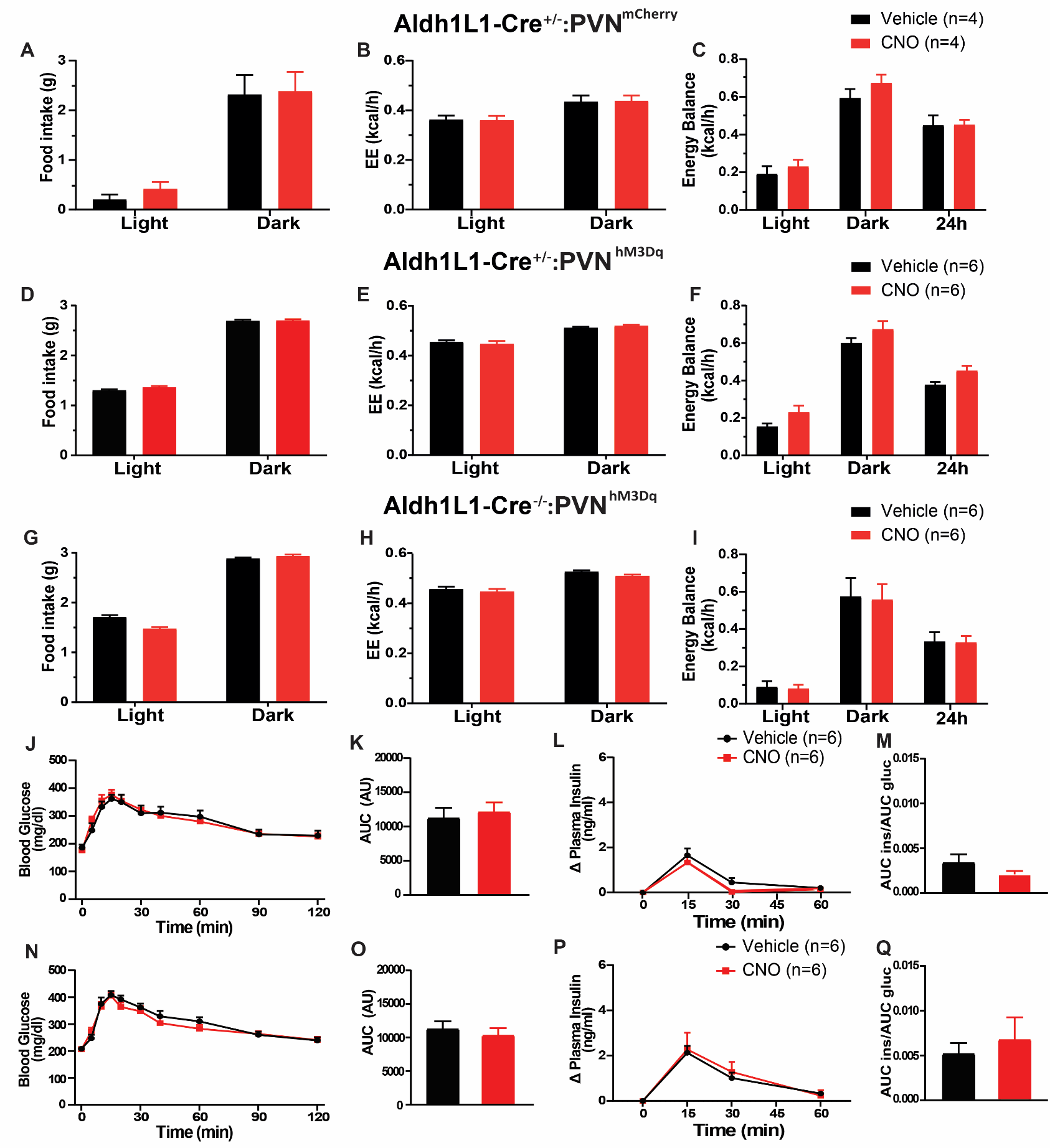

**Supplementary Figure 2. Astrocyte Gq receptor chemogenetic activation in the PVN does not modulate food intake and energy balance in lean mice. (A-I)** Food intake, energy expenditure (EE) and energy balance after Vehicle or CNO ip administration in **(A-C)** Aldh1L1-Cre^+/-^:PVN^mCherry^, **(D-F)** Aldh1L1-Cre^+/-^:PVN^hM3Dq^ and **(G-I)** Aldh1L1-Cre^-/-^:PVN^hM3Dq^ mice. **(J-Q)** Blood glucose, corresponding area under the curve (AUC), plasma insulin change and insulinogenic index after Vehicle or CNO ip administration followed by an OGTT in **(J-M)** Aldh1L1-Cre^+/-^:PVN^mCherry^ and **(N-Q)** Aldh1L1-Cre^-/-^:PVN^hM3Dq^ mice. AU refers to arbitrary units. Data are expressed as mean +/- SEM.* P<0.05. For statistical details, see table S1.

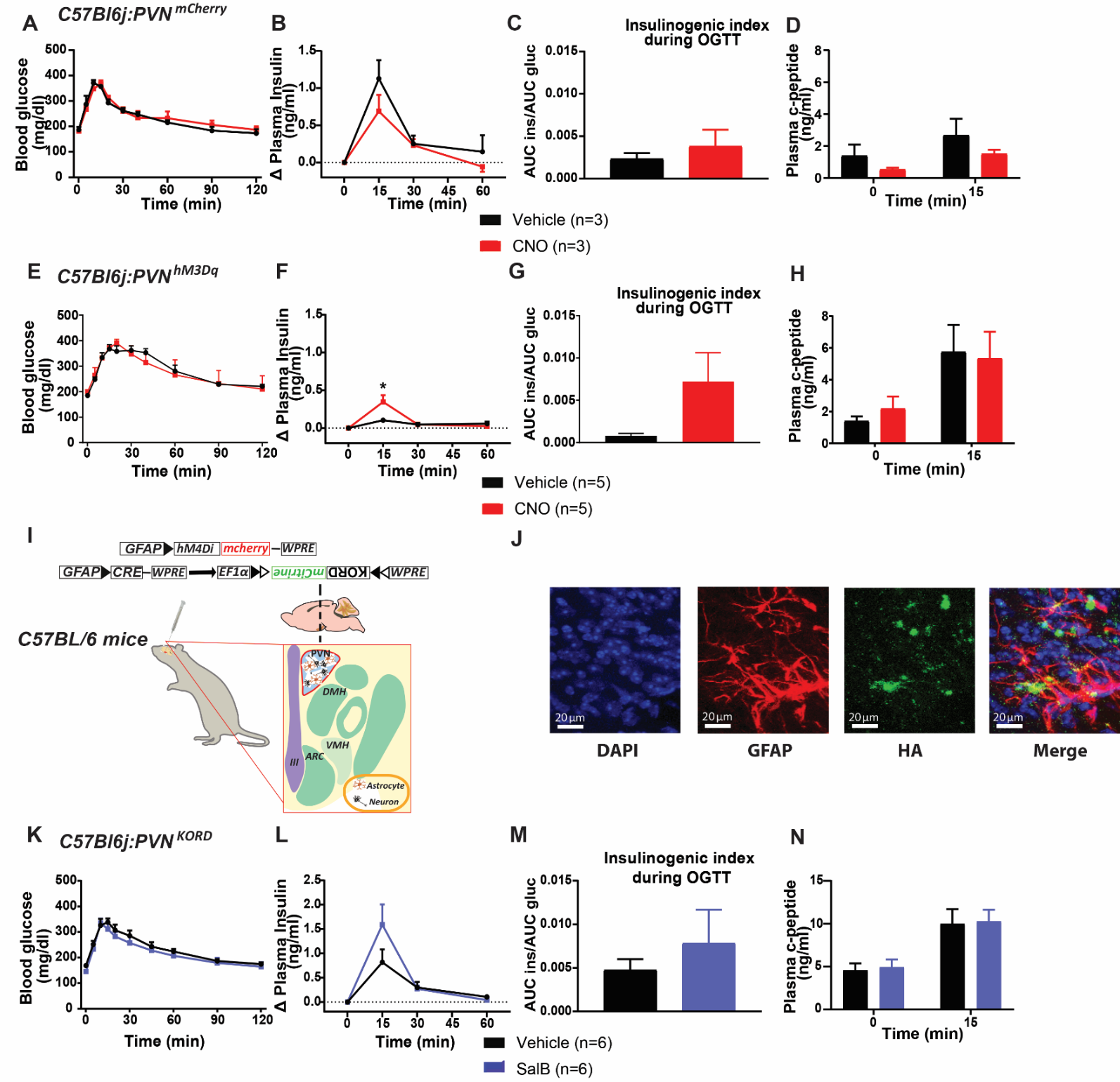

**Supplementary Figure 3. Gq receptor chemogenetic activation in GFAP-positive astrocytes in the PVN modulates glucose metabolism in lean mice. (A-H)** Blood glucose, plasma insulin change, insulinogenic index and plasma c-peptide after Vehicle or CNO ip administration followed by an OGTT in **(A-D)** C57Bl6j:PVN^mCherry^ and **(E-H)** C57Bl6j:PVN^hM3Dq^ mice. **(I)** Representative image of viral mix delivery of GFAP-Cre and Cre-KORD, to express SalB-sensitive KORD Gi receptor, or CNO-sensitive GFAP-Gi DREADD in the PVN of C57BL/6 mice. **(J)** Confocal images of double immunofluorescence of GFAP (red) and human influenza hemaglutinin (HA) tag (green) in the PVN of C57Bl6j:PVN^KORD^ mice. Cell nuclei are shown in DAPI (blue). **(K)** Blood glucose, **(L)** plasma insulin change, **(M)** insulinogenic index and **(N)** plasma c-peptide after Vehicle or SalB ip administration followed by an OGTT in C57Bl6j:PVN^KORD^ mice. Data are expressed as mean +/- SEM.* P<0.05. For statistical details, see table S1.

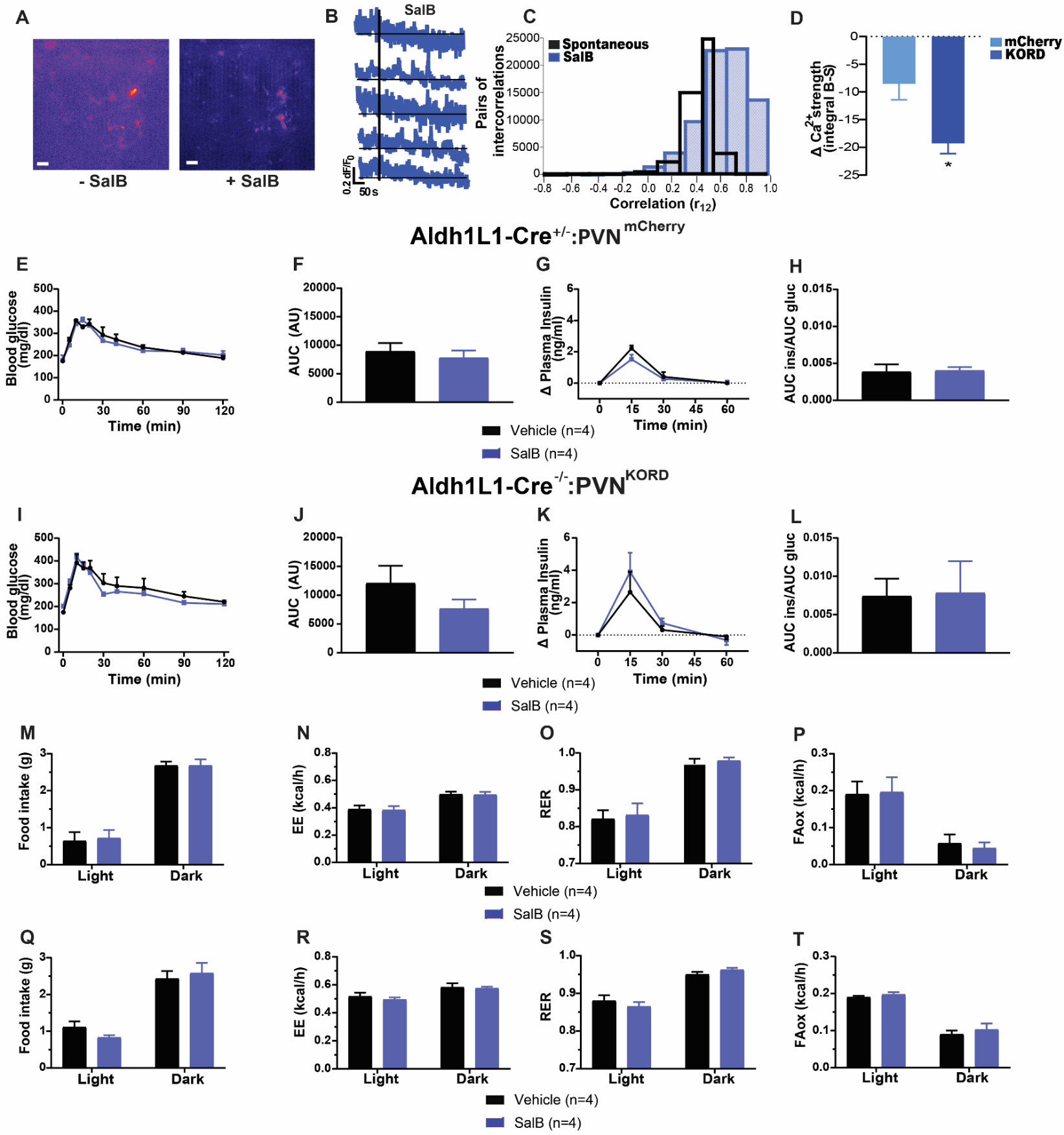

**Supplementary Figure 4. (A)** Pseudo-images, **(B)** overall Ca^2+^ strength (left) and time-course Ca^2+^ signal traces (right) of GCaMP signals, **(C)** distribution of temporal correlations between Ca^2+^ responses of all paired active domains before and after SalB bath application to PVN slices of Aldh1L1-Cre mice expressing mCherry control viral construct. **(D)** Change in overall Ca^2+^ strength after SalB bath application (integral basal-stimulation: integral B-S) to PVN slices of Aldh1L1-Cre mice expressing KORD receptor or mCherry control viral construct. Overall Ca^2+^ strength data are expressed as mean +/- SEM. Scale bar: 20µm. **(E-L)** Blood glucose, corresponding AUC, plasma insulin change and insulinogenic index after Vehicle or SalB ip administration followed by an OGTT in **(E-H)** Aldh1L1-Cre^+/-^:PVN^mCherry^ and **(I-L)** Aldh1L1-Cre^-/-^:PVN^KORD^ mice. **(M-O)** Plasma c-peptide after Vehicle or SalB ip injection followed by an OGTT in **(M)** Aldh1L1-Cre^+/-^:PVN^KORD^, **(N)** Aldh1L1-Cre^+/-^:PVN^mCherry^ and **(O)** Aldh1L1-Cre^-/-^:PVN^KORD^ mice. **(P-W)** Food intake, EE, respiratory exchange ratio (RER) and fatty acid oxidation (FAox) after Vehicle or SalB ip injection to **(P-S)** Aldh1L1-Cre^-/-^:PVN^KORD^ and **(T-W)** Aldh1L1-Cre^+/-^:PVN^mCherry^ mice. AU refers to arbitrary units. Data are expressed as mean +/- SEM.* P<0.05. For statistical details, see table S1.

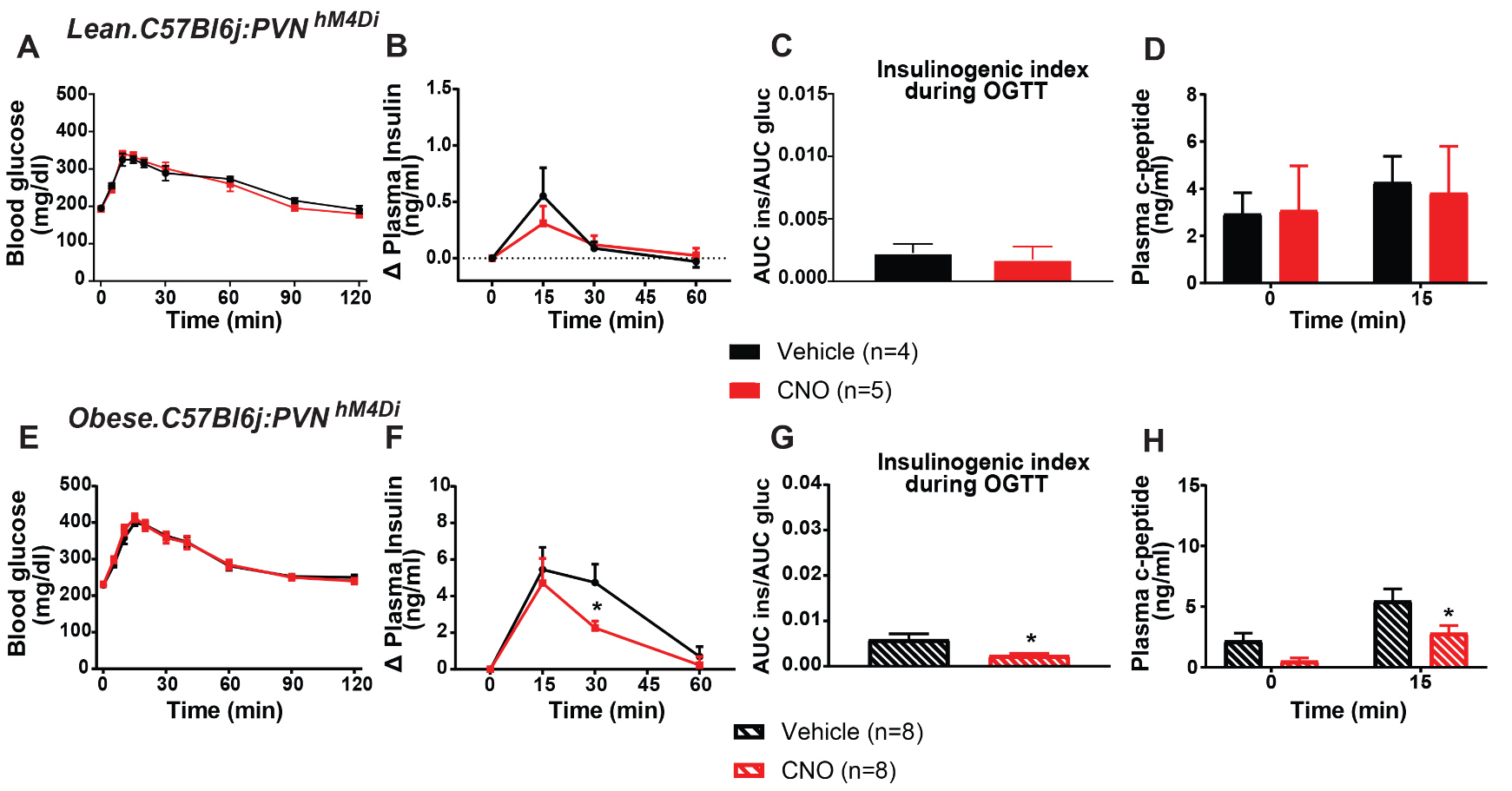

**Supplementary Figure 5. Astrocyte Gi receptor chemogenetic activation in the PVN improves glucose metabolism in obese mice. (A-H)** Blood glucose, plasma insulin change, insulinogenic index and plasma c-peptide after Vehicle or CNO ip administration followed by an OGTT in **(A-D)** lean or **(E-H)** obese C57Bl6j:PVN^hM4Di^ mice. Data are expressed as mean +/- SEM.* P<0.05. For statistical details, see table S1.

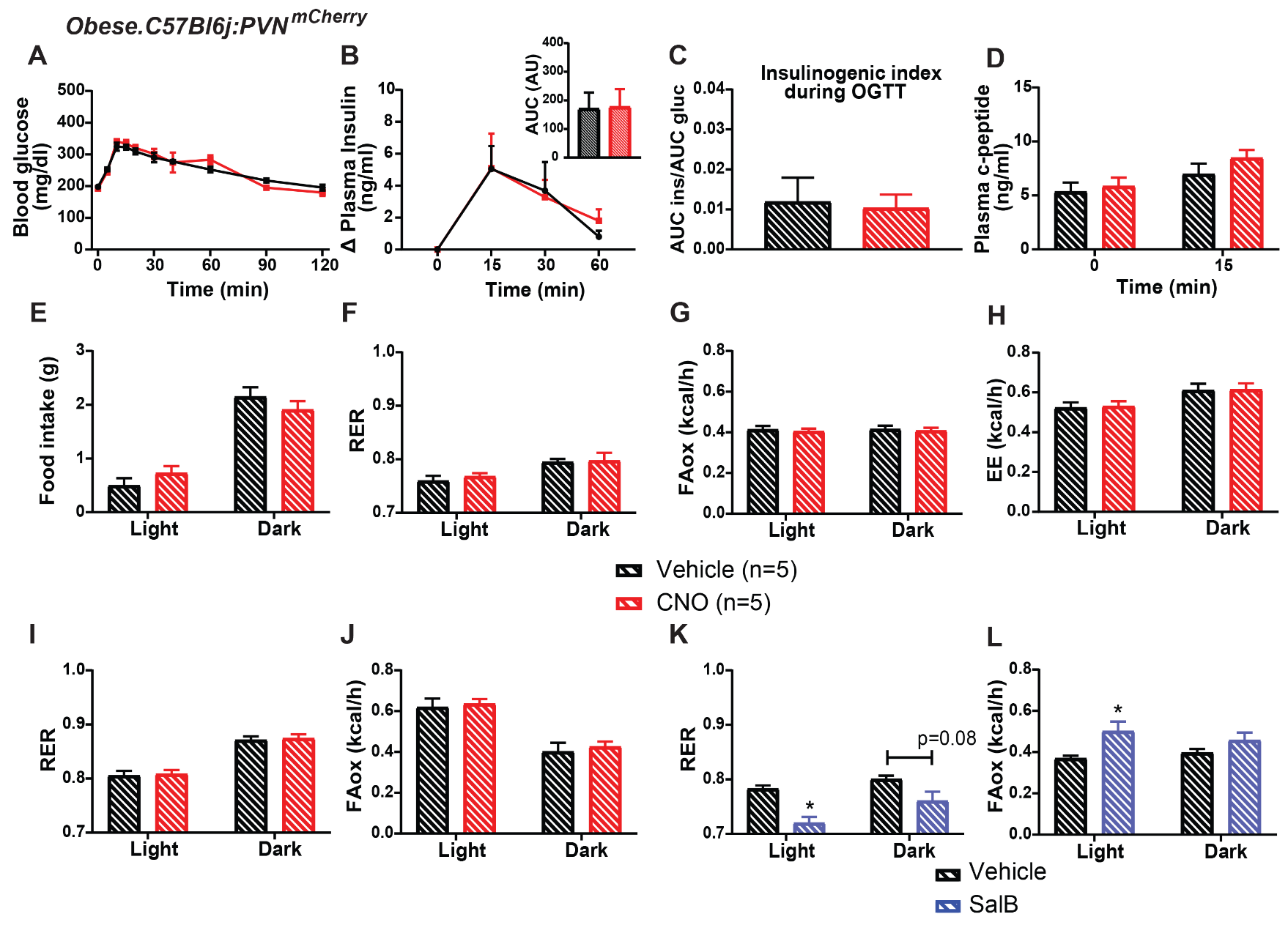
**Supplementary Figure 6. (A)** Blood glucose, **(B)** plasma insulin change and corresponding AUC (top right), **(C)** insulinogenic index and **(D)** plasma c-peptide after Vehicle or CNO ip administration followed by an OGTT in obese C57Bl6j:PVN^mCherry^ control mice. **(E)** Food intake, **(F)** RER, **(G)** FAox and **(H)** EE after Vehicle or CNO ip injection to obese C57Bl6j:PVN^mCherry^ control mice. **(I-L)** RER and FAox of obese **(I-J)** C57Bl6j:PVN^hM3Dq^ mice after Vehicle or CNO ip injection, or of obese **(K-L)** C57Bl6j:PVN^KORD^ mice after Vehicle or SalB ip administration. AU refers to arbitrary units. Data are expressed as mean +/- SEM.* P<0.05. For statistical details, see table S1.

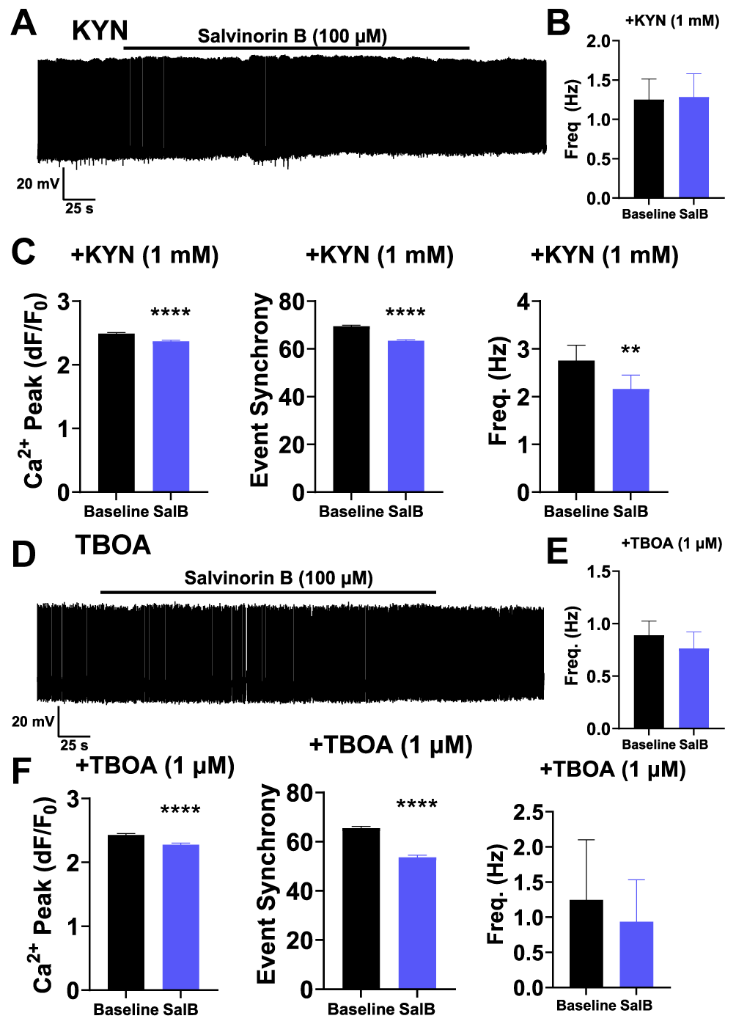

**Supplementary Figure 7. Astrocyte glutamatergic transport signaling mediates astrocyte KORD receptor-driven modulation of parvocellular neuron activity. (A)** Example trace recorded from Aldh1L1-Cre^+/-^:PVN^KORD^ mice, demonstrating the SalB effect in the presence of Kynurenic Acid (KYN, 1 mM) on parvocellular spike activity. **(B)** Parvocellular PVN neuron firing frequency in the presence of SalB and KYN in Aldh1L1-Cre^+/-^:PVN^KORD^ mice. **(C)** Summary data of astrocyte Ca^2+^ activity. Ca^2+^ strength (left), event synchrony (middle) and event frequency (right) of GCaMP signals in PVN of Aldh1L1-Cre^+/-^:PVN^KORD^ mice in the presence of SalB/KYN. **(D)** Example trace recorded from Aldh1L1-Cre^+/-^:PVN^KORD^ mice, demonstrating the SalB effect in the presence of DL-*threo*-β-Benzyloxyaspartic acid (TBOA, 1 µM) on parvocellular spike activity. **(E)** Parvocellular neuron spike frequency in the presence of TBOA and SalB in PVN slices of Aldh1L1-Cre^+/-^:PVN^KORD^ mice. **(F)** Summary data of AQuA astrocyte Ca^2+^ activity analysis. Ca^2+^ strength (left), event synchrony (middle) and event frequency (right) in the presence of TBOA/SalB. Data are expressed as mean +/- SEM. * P<0.05, ** P<0.01, *** P<0.0001. For statistical details, see table S1.

**Supplementary table 1.** Statistical analysis performed for all figures.
